# Hypoxia shapes the immune landscape in lung injury promoting inflammation persistence

**DOI:** 10.1101/2022.03.11.483935

**Authors:** Ananda S. Mirchandani, Stephen J. Jenkins, Calum C. Bain, Hannah Lawson, Patricia Coelho, Fiona Murphy, David Griffith, Ailiang Zhang, Manuel A. Sanchez-Garcia, Leila Reyes, Tyler Morrison, Simone Arienti, Pranvera Sadiku, Emily R. Watts, Rebecca. S. Dickinson, Sarah Clark, Tony Ly, David Lewis, Van Kelly, Christos Spanos, Kathryn M. Musgrave, Liam Delaney, Isla Harper, Jonathan Scott, Nicholas J. Parkinson, Anthony J. Rostron, Kenneth J Baillie, Sara Clohisey, Clare Pridans, Lara Campana, Philip Starkey-Lewis, A John Simpson, David Dockrell, Jurgen Schwarze, Nikhil Hirani, Peter J. Ratcliffe, Christopher W. Pugh, Kamil Kranc, Stuart J. Forbes, Moira K. Whyte, Sarah R. Walmsley

**Author notes:** Correspondence to: Ananda Mirchandani. Authors contributed equally (Stephen J. Jenkins, Calum C. Bain).

## Abstract

Acute Respiratory Distress Syndrome (ARDS), an often-fatal complication of pulmonary or systemic inflammation, has no cure. Hypoxemia is a defining feature, yet its impact on inflammation is often neglected. Patients with ARDS are monocytopenic early in the onset of the disease. Endotoxin or *Streptococcus pneumoniae* acute lung injury (ALI) in the context of hypoxia replicates this finding, through hypoxia-driven suppression of type I interferon signalling. This results in failed lung monocyte-derived interstitial macrophages (IM) niche expansion and unchecked neutrophilic inflammation. Administration of colony stimulating factor 1 (CSF1) rescues the monocytopenia, alters the circulating classical monocyte phenotype in hypoxic endotoxin-driven ALI and enables lung IM population expansion, thus limiting lung injury in endotoxin- and virally-induced hypoxic ALI. Hypoxia directly alters immune dynamics to the detriment of the host and manipulation of this aberrant response offers new therapeutic strategies for ARDS.

## Introduction

Acute Respiratory Distress Syndrome (ARDS) is a complication of either pulmonary or extra-pulmonary inflammation and carries a mortality rate of up to 40% ^1^. It is defined by lung infiltrates denoting lung inflammation, and systemic hypoxia, commonly requiring mechanical ventilation ^1^. Despite decades of research, effective therapies remain elusive and supportive measures are the mainstay treatment ^2^. Much of the work investigating possible therapeutic avenues has focused on modulating the inflammatory injury since its persistence is a poor prognostic indicator ^3, 4^, however the effects of systemic hypoxia per se on inflammation persistence in ARDS have not been fully explored. Macrophages have been shown to be key to driving inflammation resolution. Lung macrophages can be subdivided by their surface markers into alveolar macrophages (AM) and interstitial macrophages (IM), which inhabit distinct anatomical niches. Under homeostatic conditions, AM self-renew ^5–7^ whilst IM require recruitment of circulating monocytes ^8, 9^. Interestingly, evidence is emerging for the key role of IM in inflammation regulation ^10, 11^ and repair ^12^.

We sought to determine whether systemic hypoxia, as seen in ARDS, including severe COVID-19 disease, affected the lung monocyte-phagocyte system (MPS) and if this in turn impacted on inflammation resolution. We found ventilated ARDS patients were profoundly monocytopenic early in the disease, a phenomenon we replicated in mouse models of hypoxic acute lung injury (ALI). We further demonstrated that hypoxia fundamentally altered both monocyte phenotypes and MPS dynamics during ALI, leading to inflammation persistence, through suppression of type 1 interferon (IFN) signalling pathways. Critically, targeting this pathway with colony stimulating factor (CSF)1 corrected these hypoxia-mediated changes, driving accelerated inflammation resolution.

## Results

### ARDS is characterised by monocytopenia and an altered immune phenotype

To determine whether hypoxia is associated with alterations in the blood monocyte compartment, we obtained blood from ventilated ARDS patients early (<48 hours from diagnosis and soon after onset of hypoxic insult requiring ventilatory support) or late (>48 hours from diagnosis) and compared it to healthy controls (Table S1). Strikingly, early in the disease, whilst patients had elevated circulating leucocyte counts (Figure 1a), they had significantly lower circulating monocyte proportions (Figure 1b) and numbers (Figure 1c) compared to healthy controls. Late in the disease, circulating leucocyte numbers remained elevated compared to controls (Figure 1d) but monocyte proportions and counts were equivalent to the healthy control cohort (Figure 1e and 1f). Monocytes can be subclassified into “classical”, “intermediate” or “non-classical” cells, based on their expression of CD14 and CD16 ^13^. In early ARDS, an increase in intermediate monocytes at the expense of classical monocytes was seen in the cohort (Figure S1 a-c). Patient monocytes, irrespective of timepoint, expressed lower levels of the major histocompatibility class II (MHCII) marker, HLADR, with a concomitant increase in CD11b expression (Figure1g and 1h). This led us to question whether there were broader changes in the profile of circulating monocytes in patients with ARDS. Using a myeloid-associated 770 gene NanoString platform, we identified a transcriptional signature within ARDS monocytes (Figure 1i), with 41 genes differentially expressed compared to healthy controls (Figure 1j). Notably, 4 MHC complex genes (*HLA-DMA, HLA-DMB, HLA-DQA1, HLA-DRB3*), important for antigen-presenting function, were significantly downregulated in ARDS monocytes. Also downregulated were genes associated with monocyte adhesion and extravasation (*SELL*)^14^, trans-endothelial migration (*CD99*)^15^ and LPS signaling responses (*MAP2K4)*^16^ and *MAP3K14* ^17^. Proteomic survey of the sorted classical monocytes from healthy controls and patients with ARDS further revealed a phenotypic switch, with a significant increase in the abundance of secretory granule content within the monocytes in ARDS (Figure 1k, l). This was observed across all ARDS samples, including patients with SARS-Cov2-associated ARDS, and was associated with altered expression of known hypoxia regulated proteins including SLC2A3 ^18^, IGFR2 ^19^, PSMD4 ^20^ and FTL ^21^. Taken together these data demonstrate that ARDS affects both the number and nature of monocytes.

**Figure 1.**
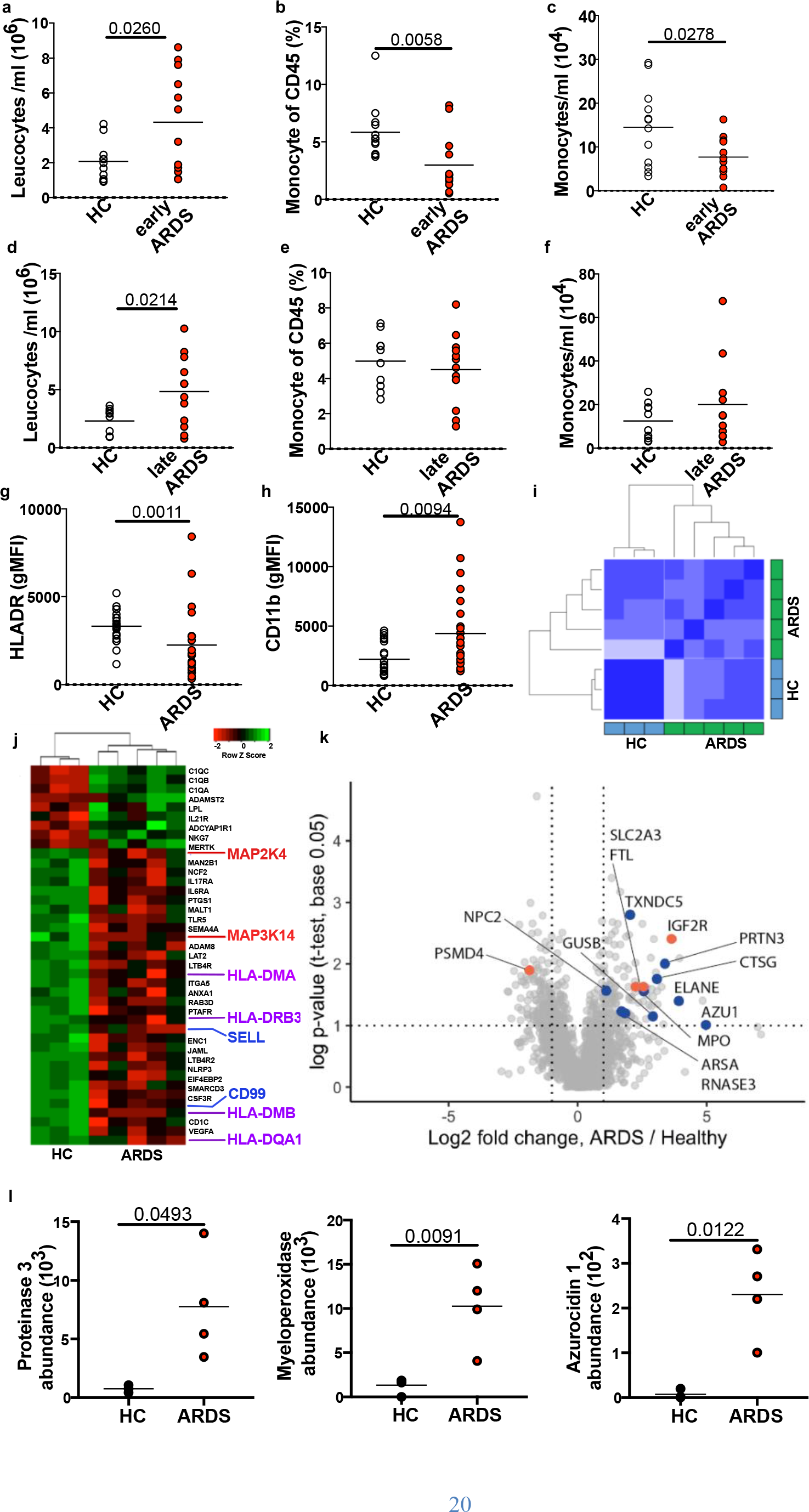
Patients with ARDS are monocytopenic early in the disease with phenotypically distinct circulating monocytes. (**a**) Blood leucocyte counts (**b**) monocyte proportions and (**c**) monocyte counts from healthy controls (HC) or patients with ARDS collected within 48 hours of diagnosis (early). (**d**) Blood leucocyte counts, (**e**) monocyte proportions and (**f**) monocyte counts from healthy controls (HC) or patients with ARDS were collected between 48 hours and 7days (late). (**g**) Monocyte HLADR and (**h**) CD11b expression in patients with ARDS compared to controls (HC n=21, ARDS n=23). (**i**) Pearson correlation and (**j**) heatmap of differentially expressed genes measured in blood monocytes from healthy control and patients with ARDS. (**k**) Volcano plot of the proteome of classical monocytes from human healthy controls (HC) and patients with ARDS (ARDS) as measured by mass spectrometry. Blue dots identify significantly-upregulated granule-associated proteins in patients with ARDS versus HC, orange dots identify a sample of proteins known to be regulated by hypoxia. (**l**) Proteinase 3, myeloperoxidase and azurocidin 1 copy numbers measured in HC and patients with ARDS. Data in **a-f,** expressed as median, **g, h**, **l** shown as mean. Each data point represents one patient. Statistical testing: **a-f, l** unpaired t-test, **g**, **h** Mann-Whitney test.

### Monocytopenia with an altered phenotype is also found in experimental ARDS

With systemic hypoxia a defining feature of ARDS, we sought to determine whether the alterations in circulating monocyte profiles in the ARDS patient cohort were consequent upon the presence of hypoxemia. Combining a model of LPS-induced ALI with reduced inspired oxygen levels (Figure 2a), allowed us to replicate the severe hypoxia and inflammatory insults that define ARDS (Figure S2a). Mice tolerated this level of hypoxia well since mice have an oxygen dissociation curve which is shifted to the right, compared to humans, allowing for improved tissue oxygenation at lower oxygen concentrations ^22^.

**Figure 2.**
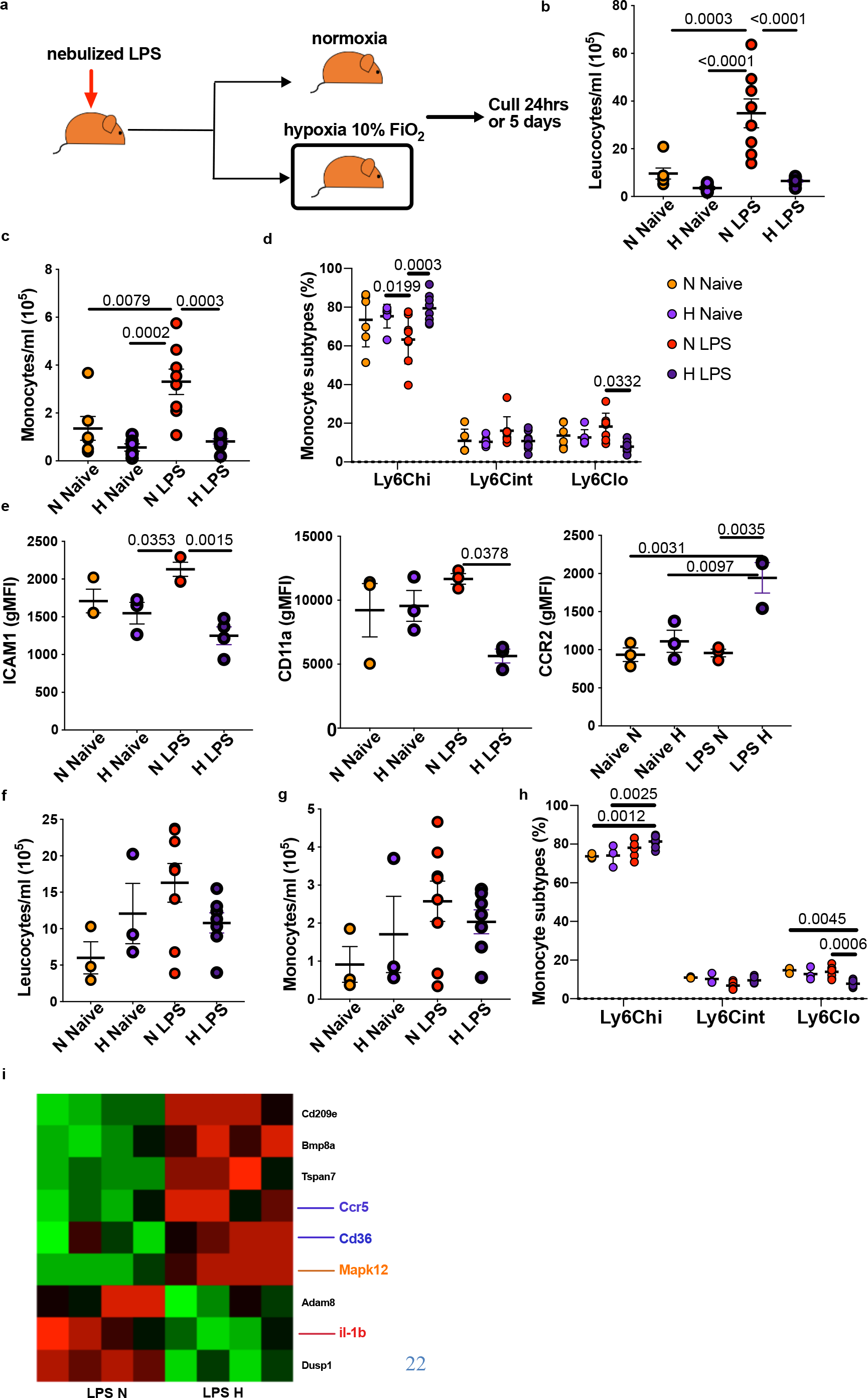
Hypoxic acute lung injury replicates early monocytopenia in mice and alters the circulating monocyte phenotype. (**a**) Experimental scheme of hypoxic or normoxic acute lung injury. (**b**) Blood cell counts and (**c**) monocyte counts in naïve or LPS-treated mice housed in normoxia (N) or hypoxia (H) for 24 hours. (**d**) Proportion of monocyte subgroups at 24. (**e**) Classical monocyte surface expression of ICAM, CD11a and CCR2 at 24hours post-LPS. (**f**) Blood cell counts, (**g**) monocyte counts and (**h**) proportions of monocyte sub-populations in naïve or LPS-treated mice housed in normoxia (N) or hypoxia (H) for 5 days post-LPS. (**i**) Differentially expressed genes in circulating classical monocytes from LPS-treated mice housed in normoxia or hypoxia for 5 days. **b-d** and **f-h** data pooled from 2 independent experiments and representative of 3-5 independent experiments, **e** data representative of 3 experiments. Each data point represents an individual mouse. Data shown as mean±SEM. Statistical testing: one-way ANOVA with Tukey’s multiple comparisons test.

The increase in circulating leukocyte counts observed following LPS challenge (Figure 2b) was abolished in hypoxic mice, with an associated reduction in circulating monocytes (Figure 2c). Moreover, monocyte populations were skewed towards a classical Ly6C^hi^ phenotype (Figure 2d), with a decrease in the non-classical Ly6C^lo^ subtype proportion.

Circulating monocytes from hypoxic mice were phenotypically distinct to those from normoxic mice with altered surface expression of adhesion molecules (ICAM1 and CD11a) and monocyte chemokine receptors (CCR2) (Figure 2e). Again, in keeping with the late ARDS human data, the number of circulating monocytes normalised by 5 days post-LPS challenge (Figure 2f and 2g), although the proportion of Ly6C^Lo^ monocytes remained contracted (Figure 2h). This was associated with persistent alterations in the transcriptome of the Ly6C^hi^ monocyte population (Figure 2i); markers of monocyte maturity (the chemokine receptor *Ccr5* and the scavenger receptor *Cd36)* ^23^ and increased *Il1b* transcript, a cytokine that has been associated with poorer outcomes in ARDS ^24^. Thus, in a mouse model of ALI, hypoxia drives alterations in monocyte number and phenotype that parallel those described in samples from patients with ARDS.

### Systemic hypoxia prevents lung interstitial macrophage compartment expansion following ALI

Given the effects of hypoxia on circulating monocyte populations, we next explored the consequence of exposure to systemic hypoxia on the immune compartment of the lung. As expected, LPS significantly increased neutrophil numbers in the lung (Figure 3a) in both hypoxia and normoxia. However, whilst LPS administration also led to a significant expansion of the macrophage (CD45^+^ Lineage^−^ CD64^bright^) compartment in normoxia, this was completely abolished in the setting of hypoxia (Figure 3b and 3c). To identify which populations accounted for this hypoxic reduction in total macrophage counts, we further characterised the CD64^bright^ compartment (Figure 3d-i). The levels of alveolar macrophages (AM), identified on the basis of their expression of CD11c and Siglec F (Figure 3d), were equivalent between hypoxia LPS challenged and normoxia LPS challenged control mice (Figure 3e). Interstitial macrophages (IM) were identified on the basis of their lack of expression of Siglec F (Figure 3d and 3f) and further subdivided based on expression of Ly6C and MHCII (Figure 3g-i). Hypoxia significantly blunted the LPS-mediated expansion of the Siglec F^−^CD64^+^ IM population (Figure 3f), with a selective loss of the Ly6C^+^ CD64^+^ but not the Ly6C^−^MHCII^+^cells (Figure 3h and 3i).

**Figure 3.**
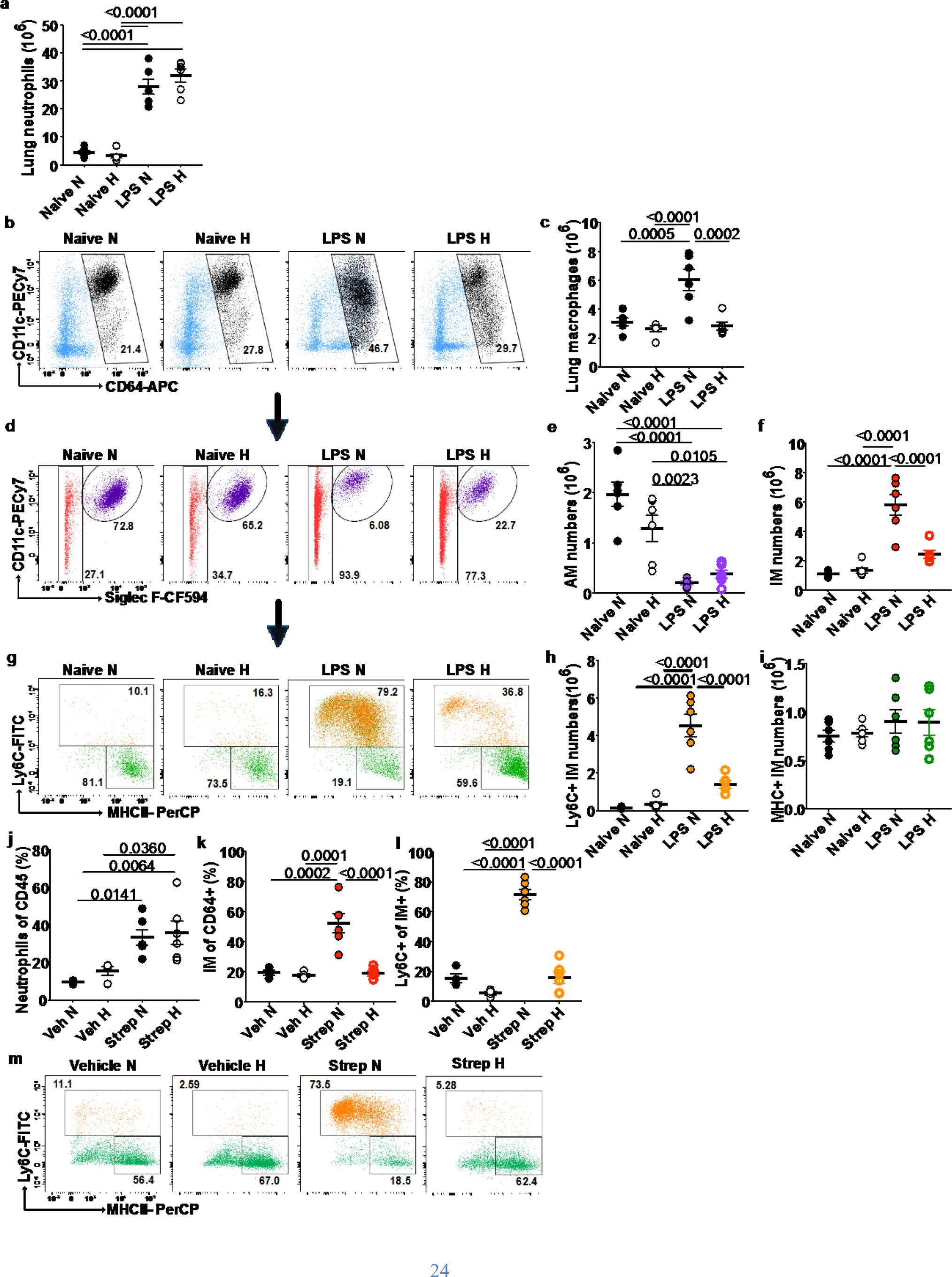
Systemic hypoxia hampers expansion of the IM niche in ALI and *Streptococcus pneumoniae* infection. (**a**) Lung neutrophils from mice treated with LPS and housed in normoxia (N) or hypoxia (H) for 24 hours. (**b**) Representative dot plot of total lung macrophages gated on Alive CD45^+^Ly6G^-^ cells and (**c**) absolute numbers of lung macrophages in naïve and LPS-treated mice housed in normoxia and hypoxia for 24 hours. (**d**) Representative dot plots of lung alveolar macrophages (AM) and interstitial macrophages (IM) gated on Alive CD45^+^Ly6G^-^CD64^bright^ and (**e**, **f**) their absolute numbers respectively. (**g**) Representative dot plots of IM sup-populations based on Ly6C and MHCII expression gated on Alive CD45^+^Ly6G^-^CD64^bright^Siglec F^-^ and (**h**, **i**) their absolute numbers. Mice were inoculated with Streptococcus pneumoniae (Strep) or vehicle (Veh) intratracheally (i.t.) and housed in normoxia (N) or hypoxia (**H**) until 24 hours post-i.t. (**j**) Lung neutrophil, (**k**) IM proportions were measured. (**l**) Ly6C+ IM sub-population absolute numbers and (M) representative dot plots of IM sup-populations based on Ly6C and MHCII expression gated on Alive CD45^+^Ly6G^-^CD64^bright^Siglec F^-^ . **c**, **e**, **f**, **h**, **i** and **j**-**m** data pooled from 2 independent experiments. **c**, **e**, **f**, **h**, **i** representative of 5 experiments. Each data point represents an individual mouse. Data shown as mean ±SEM. Statistical testing: one-way ANOVA with Tukey’s multiple comparisons test.

To complement our LPS ALI model, we examined the effects of hypoxia on monocyte/macrophage dynamics using a model of severe streptococcal pneumonia. Mice were inoculated with 10^7^ colony forming units of *D39 Streptococcus pneumoniae* and placed in either hypoxia or normoxia following a period of recovery from the anesthetic. In keeping with the results obtained in the blood in LPS-induced ALI, we saw similar changes in the blood of these mice (Figure S3a and S3b). Neutrophil recruitment was unimpaired by hypoxic conditions (Figure 3j). However, hypoxia suppressed expansion of the IM (SiglecF^−^CD64^+^) population in response to *Streptococcus pneumoniae* (Figure 3k), again with a particular loss of the Ly6C^+^ IM compartment (Figure 3l and 3m). Taken together, these data confirm that hypoxia markedly affects the MPS compartment in the lung irrespective of the nature of the inflammatory stimulus.

### Hypoxia directly alters hematopoiesis

To determine the mechanism by which hypoxia regulates circulating monocyte numbers, we measured the rate of monocyte output from bone marrow (BM) using bromodeoxyuridine (BrdU). Mice were pulsed with BrdU 12 hours after LPS nebulisation (Figure 4a) and the frequency of BrdU^+^ classical monocytes was determined 12 hours later (Figure 4b and 4c). Given that monocytes do not proliferate in circulation, any BrdU^+^ monocyte in blood must represent a recent emigrant from the BM over the preceding 12 hours^13, 25^ Consistent with our phenotypic analysis above, we found that hypoxia, irrespective of the inflammatory stimulus, caused a marked reduction in the proportion of BrdU^+^ blood monocytes (Figure 4b and 4c). Importantly, this was a selective effect on monocytes since this difference was not seen in lymphocytes (Figure S4a) and the LPS equally reduced the proportion of BrdU^+^ circulating neutrophils in hypoxic and normoxic mice (Figure S4b). This led us to question whether hypoxia was directly affecting BM hematopoiesis. Examination of the stem cell compartment (Figure 4d) at 24hrs post-LPS revealed a reduction in the number of lineage^−^Sca-1^+^Kit^+^ (LSK) cells (Figure 4e) and their sub-components, the hematopoietic progenitor cells (HPC)-1 and HPC-2 (Figure 4f and 4g) in hypoxia, independently of LPS, whilst LPS equally reduced the downstream common myeloid progenitor cells (CMP) numbers (Figure 4h).

**Figure 4.**
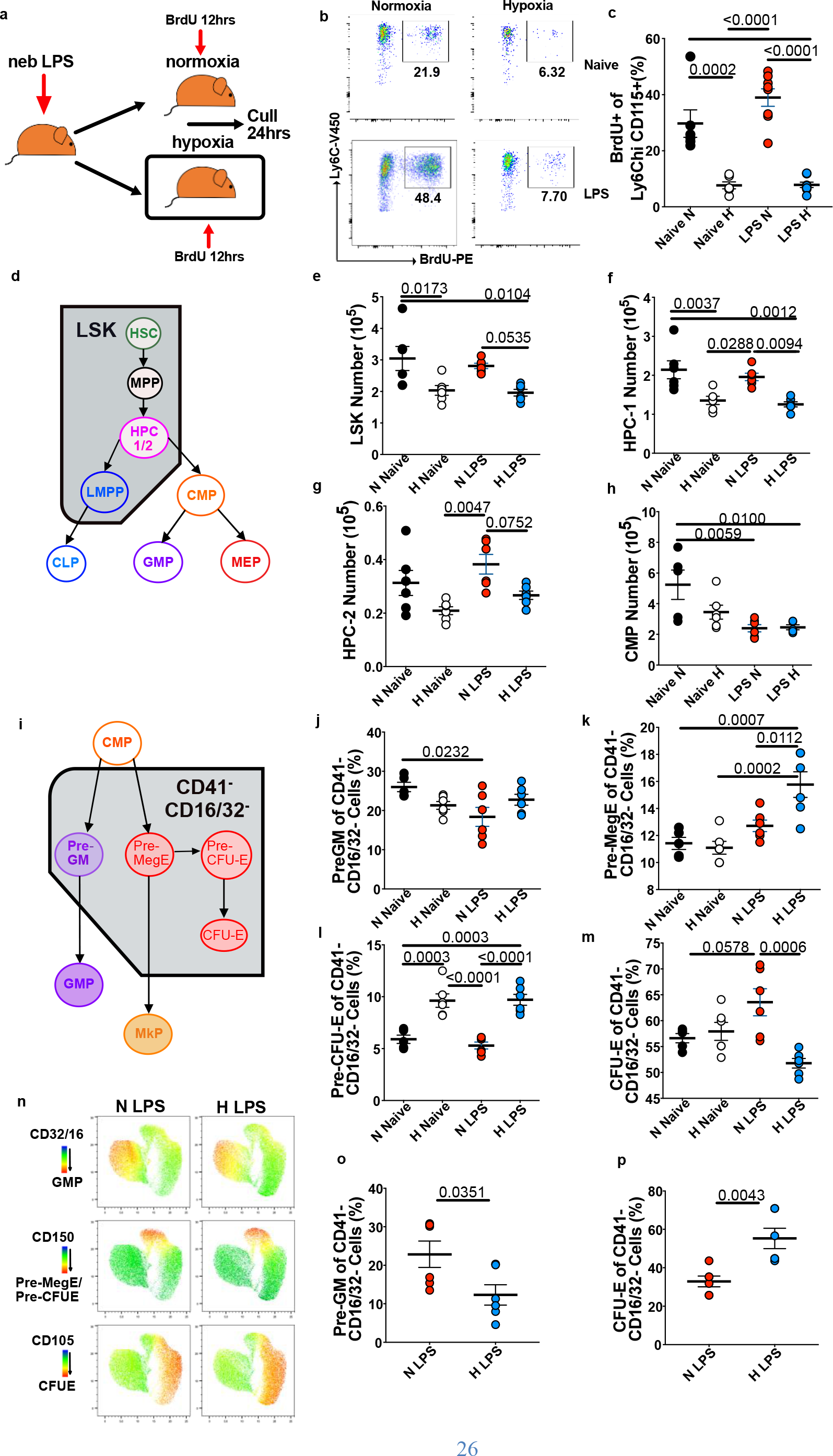
Systemic hypoxia alters bone marrow hematopoiesis towards increased erythropoiesis. (**a**) Experimental scheme for BrdU pulse in ALI. (**b**) Representative dot plots and (**c**) proportion of BrdU^+^ monocytes, gated on Alive, CD45^+^Lin^-^ (CD3/CD19/Ly6G), CD115^+^Ly6C^+^. (**d**) Schematic showing hematopoietic hierarchy with Lin^-^Sca^-^C-Kit^+^ (LSK) compartment and progenitors (HSC-hematopoietic stem cell, MPP- multipotent progenitor, HPC- hematopoietic progenitors, CMP- common myeloid progenitor, LMPP- lymphoid-primed multipotent progenitor, CLP- common lymphoid progenitor, GMP- granulocyte/monocyte progenitor, MEP-Megakaryocyte/erythrocyte progenitor). Bone marrow absolute numbers of (**e**) LSK, (**f**) hematopoietic progenitor cell (HPC)-1, (**g**) HPC-2 and (**h**) common myeloid precursor (CMP) cells. (**i**) Schematic showing erythrocytosis and monopoiesis (Pre-GM - pre-granulocyte/monocyte precursor, Pre-MegE - megakaryocyte-erythrocyte precursor, Pre-CFU-E - pre-colony forming unit erythroid, CFU-E - Colony forming unit erythroid, GMP – granulocyte-monocyte precursor, MkP- megakaryocyte precursor. (**j**) preGM, (**k**) pre-MegE, (**l**) pre-CFU-E and (**m**) CFU-E proportions were measured in bone marrow cells from mice treated with LPS and housed in normoxia (N) or hypoxia (N) for 24 hours. (**n**) Representative UMAP analysis of bone-marrow cells gated on Alive, CD45^+^Lineage^-^Sca1^-^C-Kit^+^CD41^-^CD32/16^-^ cells and summary data of proportions of (**o**) pre-GM and (**p**) CFU-E measured in bone- marrow of mice treated with LPS and housed in normoxia (N) or hypoxia (H) for 5 days. Each data point represents an individual mouse. Data shown as mean ±SEM. **a**-**c**, **e**-**m**, **o**-**p** data pooled from 2 independent experiments. Statistical testing **b**, **c**, **e-h**, **j-m** by one- way ANOVA with Tukey’s multiple comparisons test, **o-q** un-paired Student’s t-test.

Erythropoiesis in hypoxia is a highly regulated adaptive process, aimed at maintaining oxygen delivery *in vivo*. Both erythrocytes and monocytes originate from the common myeloid progenitor (CMP) ^26^ (Figure 4i and Figure S4c) leading us to question whether hypoxia also skewed hematopoiesis in favour of red blood cell production. To this end, we measured the effect of hypoxia on the CMP progeny ^26^. Whilst hypoxia did not affect the pre-granulocyte/monocyte precursors (pre-GM) population (Figure 4j) at this early timepoint, it significantly increased the relative proportion of the megakaryocyte-erythroid precursor (pre-MegE) cells in the setting of LPS-induced lung injury (Figure 4k), as well as leading to an increase in the downstream pre-colony-forming unit-erythroid precursor (pre-CFU-E) cells (Figure 4l). The CFU-E counts were diminished in the context of hypoxic ALI (Figure 4m) in keeping with increased marrow egress of red blood cells. Given that the effects of increased erythropoiesis are only measurable in the circulation after a few days, it was not surprising that the hematocrit was equivalent between the LPS-treated groups at 24 hours (Figure S4d), however we found an elevated hematocrit in hypoxic mice at day 5 post-LPS (Figure S4e). Furthermore, unbiased UMAP analysis and summary data of the CD41^-^CD32/16^-^ cells of LPS-treated mice at day 5 post-ALI demonstrated skewing in the CMP compartment within the bone-marrow was complete at that later timepoint (Figure 4n-p).

### Failed monopoiesis is a consequence of hypoxic suppression of type I interferon signalling

Human data from severely hypoxic patients with SARS-Cov2 infection has demonstrated that type I interferon responses are aberrantly blunted early in the disease ^27^ and this pathway has previously been shown to be key in regulating emergency myelopoiesis, as well as regulating erythropoiesis ^28–30^.

We therefore questioned whether hypoxia could directly regulate type 1 interferon signalling responses within the bone marrow compartment. Culture of bone marrow cells from naïve mice in hypoxia (1% FiO2) resulted in a significant blunting of type 1 IFN mediated transcriptional regulation of *Interferon Regulator Factor* (*Irf*) 8, 1 and *Ccr5* expression (Figure 5a). Subsequent analysis of the ubiquitously-expressed IFNAR receptor *in vivo* revealed a failure of LSK cells to upregulate IFNAR expression following LPS challenge in the context of hypoxia (Figure 5b). Given these findings, we postulated that hypoxic suppression of monopoiesis was consequent upon downregulation of IFNAR. Study of *Ifnar* deficient mice (*Ifnar1^−/−^*) in normoxia revealed contraction of the LSK compartment in response to LPS (Figure 5c) with skewing of haematopoiesis towards erythropoiesis (Figure 5d-f), phenocopying our findings in hypoxia. Importantly, 5 days post-LPS challenge, *Ifnar1^−/−^* mice had enhanced circulating red blood cell numbers and were significantly monocytopenic (Figure 5g and 5h). Finally, we explored whether *Ifnar^-/-^* mice replicated the myeloid lung phenotype we observed during hypoxia. Specifically, we found normal neutrophil recruitment at 24hrs to the lung (Figure 5i), equivalent AM numbers (Figure 5j), but reduced IM numbers through the failed expansion of the Ly6C^+^ IM compartment in *Ifnar1^−/−^* mice compared with controls (Figure 5k and 5l).

**Figure 5.**
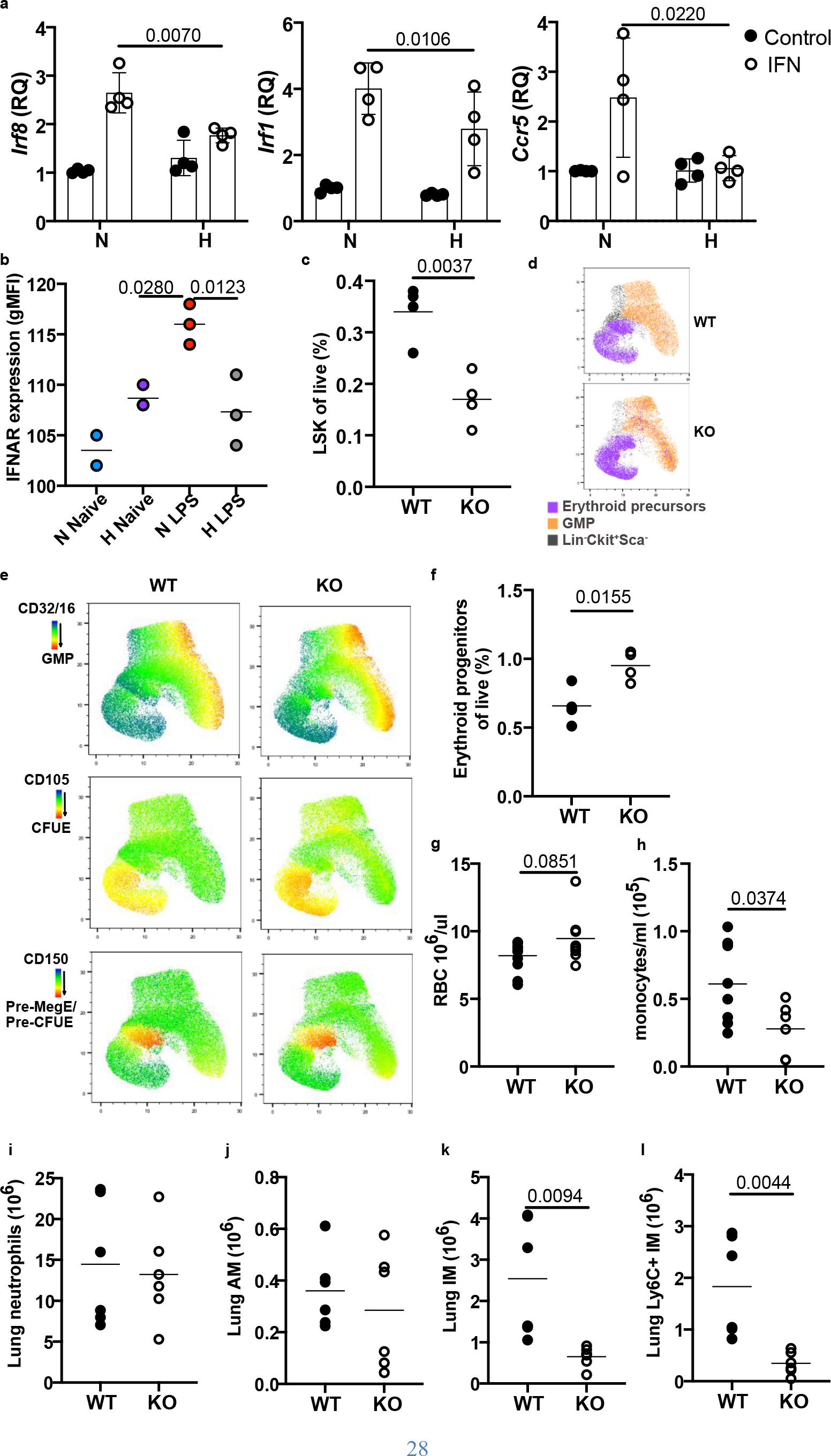
Hypoxia regulates type I interferon responses hindering lung IM expansion in response to LPS. (a) Bone marrow cells from naïve mice were cultured in normoxia (N) or hypoxia (H) for a total of 4 hours+/- interferon ß and expression of *Irf8*, *Irf1* and *Ccr5* measured by qPCR. Values were normalised to Actin ß expression. (b) Ifnar expression was measured in LSK bone marrow cells of naïve or LPS-treated mice housed in normoxia (N) or hypoxia (H) for 24hrs. (c) Proportion of LSK cells in bone marrow were measured in wild-type (WT) or *Ifnar*^-/-^ (KO) mice 24 hours post-LPS. (d) Gating of erythroid precursors and GMP in Lin^-^Ckit^+^Sca^-^ cells. (e) Representative Unbiased Uniform Manifold Approximation and Projection (UMAP) analysis of Lin^-^Ckit^+^Sca^-^ compartment demonstrating expression of CD32/16 expression (GMP marker) and CD150/CD105 (erythroid progenitor-associated markers) as per Pronk gating strategy ^26^ (f) Proportion of erythroid progenitors (Combined MEP/preCFUe/CFUe) in WT and KO bone marrow at 24hrs. (g) Red blood cell (RBC) counts in peripheral blood and (h) monocyte counts at day 5 post-LPS in WT and KO mice. Lung (i) neutrophils (j) alveolar macrophage (AM) (k) interstitial macrophage (IM) and Ly6C^+^ IM numbers at 24hrs post-LPS in WT and KO mice. All data points represent individual mice. (a) Two-way ANOVA with Šídák’s multiple comparisons post-test on two pooled experiments. (b) One-way ANOVA on 2 pooled experiments. (c,f, g-l) Unpaired Student’s t-test, (g-l) two pooled experiments.

### Monocyte recruitment failure in hypoxia is associated with lung inflammation persistence

To ascertain the longer-term effects of systemic hypoxia on the myeloid compartment for inflammation resolution, we studied the lung 5 days after LPS challenge. Strikingly, whilst the normoxic animals demonstrated very few residual neutrophils within the bronchoalveolar space 5 days after LPS challenge, mice exposed to concurrent hypoxia displayed evidence of ongoing inflammation with significant airspace neutrophilia (Figure 6a and 6b). This was accompanied by a reduction in inflammatory airspace macrophages (Figure 6c and 6d) despite recovery in the levels of circulating monocyte populations (Figure 2g). These findings were mirrored in the lung interstitium, where neutrophil numbers were broadly greater in mice housed in hypoxia (Figure 6e), and whilst AM numbers had returned to baseline (Figure 6f), the IM compartment in hypoxia remained contracted (Figure 6g) largely as a consequence of fewer Ly6C^+^ monocyte-macrophages (Figure 6h). In keeping with a profile of ongoing inflammation, the bronchoalveolar lavage (BAL) from hypoxic mice had higher levels of CXCL1 (Figure 6i), and IL-6 (Figure 6j), which in patients with ARDS has been shown to be persistently higher in non-survivors ^24^. Moreover, mice housed in hypoxia had an exaggerated weight loss profile following LPS challenge compared to their normoxic counterparts, with failure to recover to baseline by day 5 (Figure 6k). Collectively, these data demonstrate that hypoxia-induced monocytopenia is associated with persistent lung inflammation with systemic consequences that are detrimental to the host.

**Figure 6.**
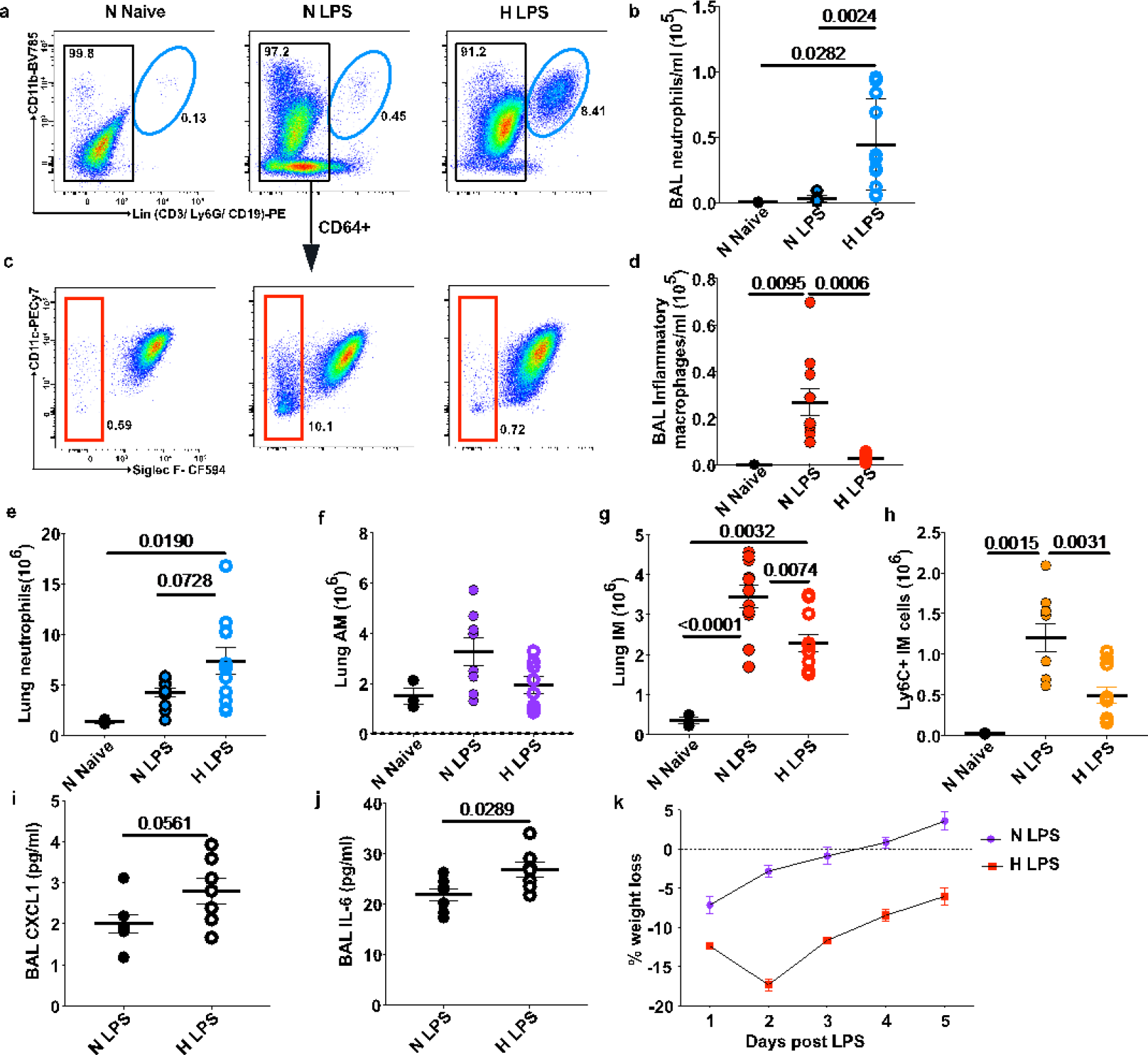
Ongoing IM expansion failure is associated with inflammation persistence in hypoxic ALI. (**a**) Representative dot plots and absolute numbers (**b**) of BAL neutrophils of mice treated with LPS and housed in normoxia (N) or hypoxia (H) for 5 days. (**c**) Representative dot plots of BAL macrophage subgroups, gated on CD64^bright^ cells, and (**d**) BAL absolute inflammatory macrophage numbers as defined by aliveCD45^+^Ly6G^-^ CD64^+^SiglecF^-^ expression. (**e**) Lung neutrophil, (**f**) alveolar macrophage (AM), (**g**) interstitial macrophage (IM) and (**h**) Ly6C+ monocyte macrophage numbers. (**i**) BAL CXCL1 and (**J**) IL-6 levels. (**k**) Weight changes following LPS in mice housed in normoxia (N) or hypoxia (H) for 5 days. Data in **b**, **d**, **e-h** pooled from 3 independent experiments, data in **i**, **j** pooled from 2 of the experiments. Each data point represents an individual mouse. Data shown as mean ±SEM. Statistical testing: **b**, **d**, **e-h** one-way ANOVA with Tukey’s multiple comparisons test. **i**, **j** by unpaired Student’s t-test.

### Systemic CSF1 treatment in hypoxia accelerates lung inflammation resolution

Finally, we tested whether specifically elevating monocyte numbers in the context of hypoxia would prove an effective therapeutic strategy and facilitate inflammation resolution. To this end, we treated LPS-challenged mice with exogenous CSF1 in the form of a CSF1-Fc fusion protein, which has been shown to prolong the half-life of CSF1 *in vivo* ^31^ (Figure 7a). As predicted, CSF1-Fc treatment vastly increased the number of monocytes in blood, with only a moderate increase in circulating neutrophil numbers (Figure 7b). CSF1-Fc treatment also increased the numbers of monocytes in the lung (Figure 7c). Whilst AM numbers were unaffected by CSF1 treatment, (Figure 7d), CSF1- Fc did lead to a marked increase in the number of CD64^+^SiglecF^−^ IM (Figure 7e). Importantly, the absolute number of neutrophils in lung tissue (Figure 7f) and BAL (Figure 7g) was reduced by CSF1-Fc treatment, despite maintained levels of CXCL1 in the BAL (Figure S5a) and elevated numbers of circulating neutrophils (Figure 7b). Mice receiving CSF1 had reduced weight loss in hypoxia post-LPS (Figure S5b). Furthermore, CSF1 treatment also reduced the IgM levels in BAL fluid (Figure 7h) in keeping with reduced alveolar inflammation and vascular leak.

**Figure 7.**
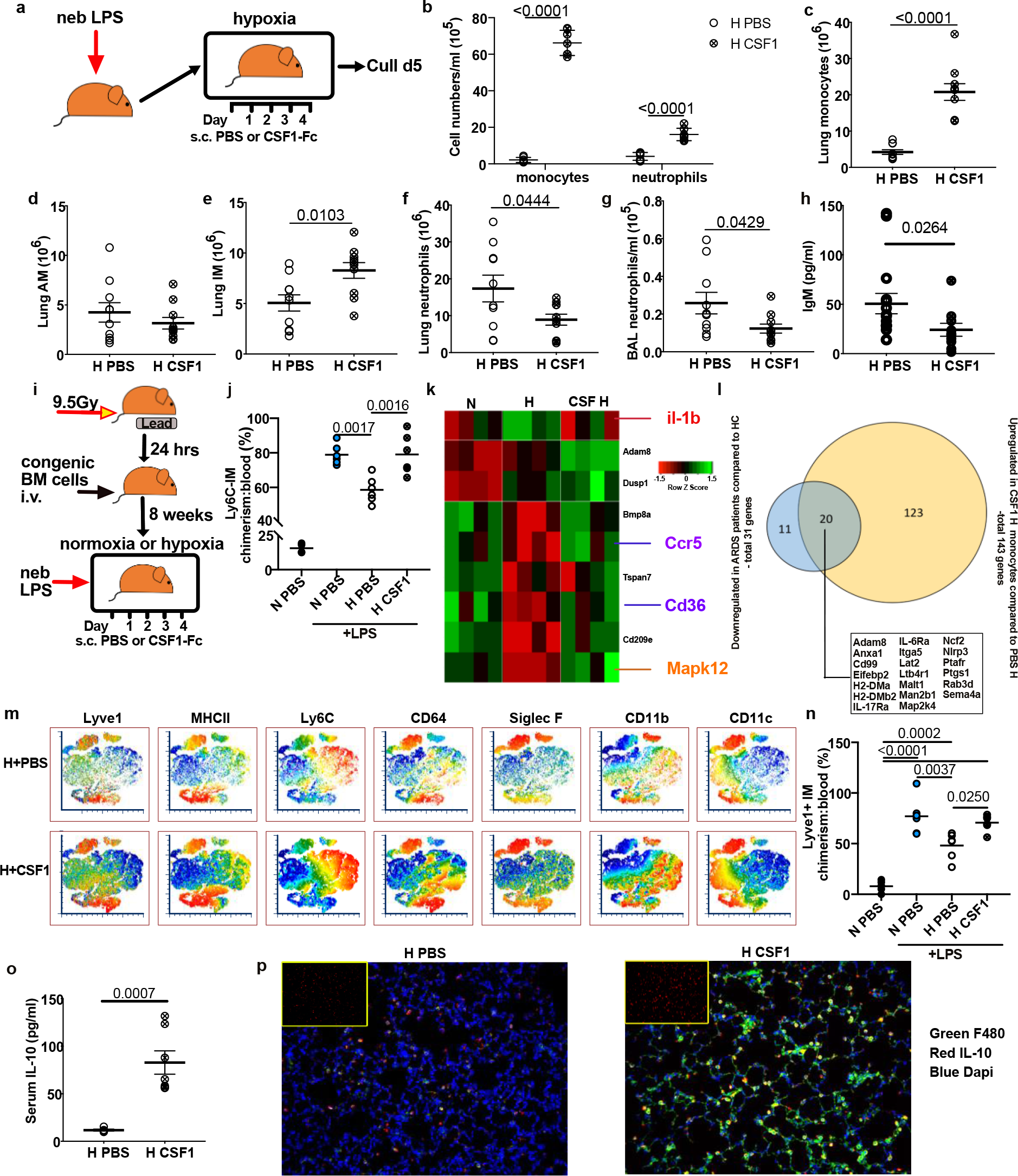
CSF1 rescues the hypoxic monocytopenia driving inflammation resolution. (**a**) Experimental scheme for CSF1-Fc treatment in hypoxic ALI. (**b**) Blood monocyte and neutrophil numbers in PBS and CSF1-Fc-treated mice housed in hypoxia (H). (**c**) Lung monocyte (**d**) alveolar macrophage (AM), (**e**) interstitial macrophage (IM), (**f**) lung neutrophil and (**g**) bronchoalveolar (BAL) BAL neutrophil numbers in hypoxic ALI PBS- or CSF1-treated mice. (**h**) BAL IgM as measured by ELISA in hypoxic ALI PBS- or CSF1-treated mice. (**i**) Schematic of tissue-protected chimera and LPS-induced ALI model. (**j**) Lung Ly6C-IM:blood monocyte chimerism in mice with or without LPS-induced ALI, treated with PBS or CSF1 and housed in normoxia (N) or hypoxia (H). (**k**) Classical blood monocyte heatmap of differentially expressed genes in LPS-treated mice at day 5 (**l**) Venn diagram showing overlap of differentially downregulated genes in patients with ARDS, that are upregulated by CSF1 treatment compared to PBS alone (all genes fold change ≥1, except H2-DMa, H2-DMb2, IL-17Ra, Nlrp3 which were upregulated by CSF1 >0.5 fold-change, p<0.05). (**m**) Representative images of tsne analysis of AliveCD45^+^Ly6G^-^CD11b^+^ and/or CD45^+^Ly6G^-^ CD11c^+^ lung cells. (**n**) Lyve1^+^IM: blood monocyte chimerism . (**o**) serum IL-10 levels were measured in mice with LPS-induced ALI, housed in hypoxia for 5 days, and treated with CSF1-Fc or PBS. (**p**) Representative Immunofluorescence of lung sections from mice with LPS-induced ALI, housed in hypoxia and treated with PBS or CSF1. IL-10 alone in inset image with merged triple stain in main. Data shown as mean ±SEM with each data point representing an individual mouse. Statistics **b**-**h**, **o** unpaired Students t-test. **j**, **n** One-way ANOVA with Tukeys post-test, **h** two-tailed Mann-Whitney following D’Agostino & Pearson normality test**. b-h** data pooled from 3 independent experiments, **j**, **n**, **o** data pooled from 2 independent experiments.

To extend our *in vivo* observations to a model of ARDS driven by virally-induced epithelial injury, mice were challenged with Influenza A in the context of hypoxia (Figure S5c). In keeping with our observations in sterile lung injury models, CSF1-Fc treatment increased the number of monocytes recruited into the lung in hypoxic PR8-infected mice (Figure S5d), improving physiological outcomes (Figure S5e). This was associated with significant reductions in markers of lung injury (Figure S5f) and cellular damage (lactate dehydrogenase activity) (Figure S5g).

In order to dissect the mechanism whereby CSF1-Fc accelerated inflammation resolution, we used lung-protected bone marrow chimeric mice (Figure 7i) to track the ontogeny of the expanded IM compartment in LPS-treated mice housed in either hypoxia or normoxia. CD103^+^ dendritic cells (cDC1), as well as CD11b^+^ DC (cDC2) were also fate-mapped using this system (Figure S5h and S5i). Consistent with previous work ^32^, we found the non-host chimerism of IM from unchallenged chimeric mice maintained in normoxic conditions to be around ∼20% when normalised to that of blood monocytes, indicative of a slow rate of replenishment of these cells under normal conditions. Administration of LPS in normoxia led to a dramatic increase in chimerism, in keeping with enhanced recruitment of bone marrow-derived cells into the IM compartment. Importantly, in hypoxia, administration of exogenous CSF1-Fc rescued the failed expansion of the IM compartment (Figure 7j) through recruitment of circulating monocytes as evidenced by increased non-host chimerism in the CSF1-Fc treated mice.

Whilst it was clear that CSF1 increased the numbers of circulating monocytes, and in turn, replenished the lung IM, we wanted to establish whether CSF1 therapy also rescued the phenotypic shift seen in hypoxic circulating monocyte populations. CSF1-Fc treatment normalised the expression of the genes we previously reported to be suppressed by systemic hypoxic exposure, including the type I IFN-associated gene *Ccr5*, without enhancing the expression of the pro-inflammatory *Il1b* (Figure 7k and Figure S5j).

Interestingly, CSF1 therapy significantly induced a number of genes we had observed to be suppressed in ARDS patient samples (Figure 7l). These included key extravasation molecule genes such as *Itga5, CD99* and *Sell* and anti-inflammatory genes such as *Anxa1* ^33^. Importantly, these CSF1-treated monocytes gave rise to phenotypically distinct IM as demonstrated by unbiased analysis of the lung MPS compartment, consequent upon expansion of the CD64^bright^ MHCII^−^ fraction. Of note, many of these MHCII^−^ cells expressed high levels of Lyve-1 (Figure 7m), making them phenotypically identical to repair-associated macrophages that arise during bleomycin-induced lung injury ^12^. Fate mapping revealed expansion of these MHC-Lyve1+IM cells to be consequent upon circulating monocyte recruitment to the lung (Figure 7n).

To delineate the mechanism by which CSF-1 mediated expansion of the IM population facilitates inflammation resolution and clearance of neutrophils recruited to the lungs we looked for altered expression of known mediators of efferocytosis ^34^. With evidence of elevated levels of IL-10 in the serum of CSF1-treated mice (Figure 7o) we sought to determine whether CSF1 treatment led to an increase in IL-10+ IM in the lung. Using immunofluorescence, we confirmed CSF1 treatment greatly increased the numbers of IL- 10 expressing interstitial F480+ cells in the lung (Figure 7p). Thus, CSF1 not only altered monocyte-macrophage numbers in the lung, but also drove the resolution of the dysfunctional neutrophilic inflammation generated by hypoxia and promoted restoration of normal tissue homeostasis.

Taken together, these results underpin the importance of monocyte recruitment into the lung during hypoxic lung inflammation in order to drive resolution, as well as the therapeutic potential of CSF1 as a treatment in non-resolving ARDS

## Discussion

Hypoxia and alveolar inflammation are the two defining features of ARDS and inflammation persistence in ARDS is a well-known poor prognostic feature ^35, 36^ We have previously demonstrated how hypoxia negatively affects the host’s capacity overcome bacterial ALI ^37^ and this study expands on those findings to define the impact of hypoxia on the host’s capacity to drive inflammation resolution.

Monocytes are professional phagocytes ^38^, and as such, are key mediators of restoration of homeostasis ^39^. We show that, despite the heterogeneity in the aetiology of disease, patients with early ARDS, including those infected with SARS-CoV-2, are monocytopenic and display a defective functional profile with markedly reduced HLADR expression and increased secretory granule content. Modelling hypoxic ALI with LPS and *Streptococcus pneumoniae* challenge in the mouse, we equally observed an early monocytopenia, which resulted in a failure to expand the IM compartment in the lung, as confirmed by lung- protected chimera studies. In vivo BrdU pulse chase experiments and analysis of bone marrow progenitors subsequently identified an important consequence of systemic hypoxia to be contraction of the stem cell compartment with skewing towards erythropoiesis.

In the context of prioritising the preservation of tissue oxygen-delivery, the skewing towards erythropoiesis makes physiological sense, for example in adaptation to altitude ^40^ where monocytopenia has been observed as early as 1969 ^41^. However, when engagement of an effective innate immune response is also required, our data would suggest that these hypoxia-induced changes have consequences for inflammation resolution with persistence of neutrophilic inflammation. This is clinically important as evidenced most recently by the current COVID-19 pandemic, in which disordered innate and adaptive host responses co-exist with severe systemic hypoxia ^42, 43^.

Monocytes appear to be highly plastic and their phenotype has been shown to be determined before their release into the bloodstream. Yanez et al demonstrated that the monocyte phenotype generated in the bone marrow is dependent on the nature of the insult driving the emergency myelopoiesis, hence a TLR4 ligand drives “neutrophil”-like monocyte production, whereas a TLR9 ligand generates a more “DC”-like cell ^44^. This balance, however is in the context of normoxia. Interestingly, the presence of a “bacterial” signature, with suppressed type I IFN responses has been shown in patients severely unwell with influenza ^45^ whilst ifnar-/- monocytes have increased expression of granule-associated genes ^46^. Importantly, data from our ARDS patients demonstrate that this “bacterial” phenotype was present in both bacterial- and viral-induced ARDS. Furthermore, our *in vivo* data show that hypoxia per se, inhibits type I interferon responses in the bone marrow, whilst leading to in enhanced lung injury and impaired inflammation resolution. We therefore propose that, the systemic hypoxia that defines severe lung injury and ARDS, is not merely a marker of disease severity, but a key orchestrator shaping the immune responses in these conditions. Furthermore, hypoxic-driven inhibition of type I IFN may partly explain the association of hypoxia with poorer outcomes in viral-induced ALI, such as severe COVID-19 disease ^47^. A limitation of our current work and an area of future research focus remains the elucidation of the mechanisms whereby hypoxia inhibits inflammation-driven IFNAR upregulation, and the effects of hypoxia on this pathway beyond the bone marrow.

There are no disease-modifying therapies available for ARDS, and in severe COVID-19 disease, most therapies are aimed at dampening the pro-inflammatory responses in the host. By using CSF1 as a therapy we propose to alter the immune responses at their origin, enhancing the pro-resolution capacity of the monocytes released in the blood stream. The therapeutic expansion of monocyte-derived macrophages in the lung may appear to be counterintuitive given data in which the presence of ‘activated’ monocytes are associated with poor outcomes following coinfections ^43^. Moreover, proinflammatory monocytes have been also linked to the cytokine storm observed in patients with severe COVID-19 disease ^48^. The finding that CSF1-Fc, not only increases the number of monocytes and IM in the lung, but also alters their phenotype, helps explain this discrepancy. At a transcriptional level, CSF1 treatment broadly restored the signature of circulating monocytes from hypoxia challenged mice to that observed in a normoxic state. It also upregulated 60% of the genes that were suppressed in the human ARDS monocyte populations. These phenotypic differences occurred in the CSF1-Fc-treated group despite the IM developing in a highly inflamed and hypoxic environment. This is particularly interesting given that macrophages demonstrate great plasticity and the niche they inhabit is key to their functional imprinting, as reviewed recently ^49^.

In our system, CSF1 treatment led to a vast increase in IL-10^+^ IM with a concomitant increase in systemic IL-10. Macrophages are known to release IL-10 upon efferocytosis, driving resolution of inflammation ^34^. Furthermore, IL-10-producing IM have recently been shown to be key regulators of both allergic ^9^ and endotoxin-mediated lung injury ^11^. CSF1 treatment also led to an increase in the number of Lyve1^+^ IM, a marker associated with a subgroup of macrophages that have been shown to be key to tissue repair in various disease contexts ^50, 51^, including in the lung ^12^. These findings strengthen the case of the therapeutic potential of manipulating the CSF1 axis in human ARDS. However, if CSF1, or indeed any MPS cell therapy, is to be considered as a therapeutic in ARDS, ensuring that the cells induced, or delivered, retain their pro-repair phenotype within the highly inflamed niche they find themselves in, is fundamental. Further work in this area is therefore warranted.

Taken together, our findings demonstrate the importance of hypoxia as an active participant in defining the immune landscape, and therefore outcomes, in acute lung injury. These findings, compounded with the effect of CSF1-Fc on accelerating inflammation resolution and reducing lung injury, underscores the therapeutic potential of CSF1 as a novel treatment in ARDS.

## Acknowledgments

We are grateful to Prof D Hume for providing the CSF1-Fc used in these experiments. We are grateful to Thomson Bioinformatics, 27 Strathalmond Road, Edinburgh, United Kingdom, for analysing the NanoString data. Flow cytometry data were generated with support from the QMRI Flow Cytometry and Cell Sorting Facility, University of Edinburgh. We are grateful to the Royal Infirmary of Edinburgh Critical Care Research Team for their assistance in recruiting, consenting and obtaining samples from patients with ARDS. This work was funded by a Wellcome Trust Senior Clinical fellowship awarded to S.R.W. (098516 and 209220), Wellcome Trust Post-doctoral Training Clinical Fellowship (110086) and a Wellcome Trust iTPA grant (PIII052) awarded to A.S.M. and was partly funded by UKRI/NIHR through the UK Coronavirus Immunology Consortium (UK-CIC). C.C.B holds a Sir Henry Dale Fellowship jointly funded by the Wellcome Trust and the Royal Society (Grant Number 206234/Z/17/Z).

## Author contributions

A.S.M, S.J.J, C.C.B, H.L, K.K, M.K.W, and S.R.W. conceived and designed the experiments, A.S.M, C.C.B, H.L, P.C, F.M, M.A. S-G, LR, TM, SA, R.D, E.R.W, L.D, D.G, I.H, J.S, A.Z, L.C, D.L, V.K, C.S, N.J.P and P.S-L performed the experiments. A.S.M, C.C.B, H.L, T.L analysed the data. A.S.M, C.C.B, S.J.J, H.L, K.K, S.J.F, C.P, P.J.R. M.K.W and S.R.W. interpreted the data. S.C, K.M.M, J.S, A.J.R, A.J.S, D.D facilitated obtaining patient samples. A.S.M S.J.J. N.H. M.K.W. and S.R.W. helped obtain funding. K.B and C.L provided reagents. J.S. provided ifnar^-/-^ KO mice. A.S.M, S.J.J, C.C.B, H.L, K.K, C.W.P, S.J.F, M.K.W and S.R.W. wrote the manuscript.

## Methods

### RESOURCE AVAILABILITY

#### Lead contact

Further information and requests for resources and reagents should be directed to and will be fulfilled by the Lead Contact, Ananda Mirchandani

#### Materials Availability

This study did not generate new unique reagents.

#### Data and Code Availability

Analyses, including the drawing of heatmaps and volcano plots were carried out in R using the package ggplot2 (https://cran.r-project.org/web/packages/ggplot2/index.html). Analysis of datasets was carried out by Thomson Bioinformatics, Edinburgh, UK. **EXPERIMENTAL MODEL AND SUBJECT DETAILS**

#### Human healthy control blood donors

Patients with ARDS were recruited and informed consent obtained directly or by proxy under the “META-CYTE” study (17/SS/0136/AM01) and “ARDS-NEUT” study (20/SS/0002) as approved by the Scotland A Research Ethics Committee. Samples were also obtained under the “Effects of Critical Illness on the Innate Immune System” study as approved by Health Research Authority (REC number 18/NE/0036).

All healthy participants gave written informed consent in accordance with the Declaration of Helsinki principles, with AMREC approval for the study of healthy human volunteers through the MRC / University of Edinburgh Centre for Inflammation Research blood resource (15-HV-013).

Up to 20-40mls of whole blood was collected into citrate tubes and up to 10 million cells were stained for flow cytometry assessment and sorting. Briefly, the whole blood was treated with red cell lysis buffer (Invitrogen) and cells counted prior to staining for flow cytometry. Cells were incubated with anti-CD16/32 Fc-block (2:50) for 30 minutes followed by staining for 30 minutes with antibodies (see Table 1) followed by a wash with FACS buffer (PBS+2% Fetal calf serum (FCS)). Dapi (1:1000) was added prior to flow cytometry to determine live cells. Monocytes were identified as Singles Dapi^-^CD45^+^non-granulocyte Lin(CD3/CD56/CD19+/-CD66b)^-^ HLADR^+^ CD14^+^ and/or CD16^+^ cells.

Samples obtained from April 2020 were fixed prior to acquisition given the potential for SARS-Cov2 dissemination. Briefly, 1uL of zombie Aqua fixable viability dye (stock 1:20 dilution) was added to 100uL of whole blood for 15 minutes at room temperature in the dark. 2uL of FcBlock was added for a further 30 minutes, on ice. Samples were then stained as above and fixed/ lysed using BD FACS Lyse for 10 minutes at room temperature. The sample was then resuspended in 300uL of FACS buffer and 50uL of Countbright beads added (Thermofisher) prior to acquisition.

#### Animals

Male C57/BL6 mice aged 8-15 weeks were purchased from Envigo or Charles River.

*Ifnar^-/-^* (*ifnar ^tm/agt^*) mice were obtained from J.S. who purchased them originally from the Jackson laboratory.

Animal experiments were conducted in accordance with the UK Home Office Animals (Scientific Procedures) Act of 1986 with local ethical approval.

#### Mouse LPS ALI Model

Mice were treated with nebulized LPS (3mg), and were then housed in normoxia or hypoxia (10% O2) for up to 5 days. Mice were treated with Colony Stimulating Factor (CSF)-1-Fc (kind gift from Prof D Hume) by subcutaneous injection (0.75μg/g/mouse) from day 1 to 4 post-LPS, prior to cull on day 5.

#### D39 Strep. Pneumoniae infection

Mice were anesthetized and 10^7^ colony forming units (or vehicle) were delivered in 50uL PBS via intratracheal intubation. Following reversal of anesthetic and a period of recovery, mice remained in normoxia, or were placed in hypoxia.

#### Influenza A (PR8) virally-induced ALI model

Mice were lightly anesthetized using isofluorane and 20 plaque-forming units (p.f.u.) of PR8 Influenza A virus in Dulbecco’s Modified Eagle’s Media were inoculated intranasally. After 1 hour of recovery time, mice were placed in hypoxia for 48 hours. Subcutaneous PBS or CSF1-Fc injections (as above) at 12 hours and 36 hours were administered. Sickness scores were determined using methods as described previously ^37^.

#### Lung and alveolar cell sampling

Mice were culled with an overdose of intraperitoneal anesthetic (Euthetal) followed by blood collection from the inferior vena cava. Alveolar leukocytes were collected by bronchoalveolar lavage (BAL), then mice were perfused lightly with PBS through the heart, prior to harvest of lung tissue. On occasion, lower limbs were harvested for bone marrow leukocyte assessment (see below).

Tissue leukocytes were extracted from surgically dissociated lung tissue by enzymatic digestion with 2ml enzyme mix (RPMI with 0.625 mg/ml collagenase D (Roche), 0.85 mg/ml collagenase V (Sigma-Aldrich), 1 mg/ml dispase (Gibco, Invitrogen) and 30 U/ml DNase (Roche Diagnostics GmbH) for 45 minutes at 37°C in a shaking incubator. The digest material was passed through a 100μM cell strainer with the addition of FACS buffer (PBS with 0.5% BSA/2% fetal calf serum and 0.02mM EDTA). Cell pellets were treated with red cell lysis buffer (Sigma) and washed in FACS buffer. The resulting cell suspension was subsequently passed through a 40μm strainer before cell counting using a Casey TT counter (Roche). Single cell suspensions (5 million cells/sample) were then stained for flow cytometry. BAL samples were counted prior to staining for flow cytometry. ***Blood and bone marrow sampling***

Mouse blood and bone marrow were treated with red blood cell lysis buffer (Biolegend) prior to counting and staining for flow cytometry (see Table 1).

Hematopoietic cell assessment was performed using both hind legs that were crushed using a pestle and mortar until a homogenous cell suspension was achieved or flushed though using a 32G needle. Cells were collected in cold FACS buffer and filtered through a 70 um nylon strainer (BD Falcon, 352340). Cells were treated with RBC lysis buffer (Biolegend) prior to staining

#### Tissue protected chimeras

6–8-week old C57BL/6J CD45.1^+^CD45.2^+^ mice were anesthetized and were irradiated with a single dose of 9.5 Gy γ-irradiation while all but the hind legs and lower abdomen were protected by a 2 inch lead shield. The following day, the mice received 2-5 × 10^6^ BM cells from CD45.2^+^ C57BL/6J by iv injection.

### METHOD DETAILS

#### Flow Cytometry

Mouse cells were treated with α-CD16/32 Fc block (e-bioscience) (1:100) prior to staining with antibodies (see supplementary table 1). Relevant fluorescence minus one (FMO) samples were used as controls. Zombie Aqua fixable viability dye (Biolegend) was used prior to Fc block to exclude dead cells from digest samples or Dapi for single cell suspensions.

Cells were acquired on the LSRFortessa (Becton Dickinson) or sorted on an Aria II or Fusion machine (Becton Dickinson). Compensation was performed using BD FACSDiva software and data analyzed in FlowJo version 10 or FCS Express 7 for tsne analysis.

| **Antibody** | **Clone** | **Catalogue** | **Lot** | **Fluorophore** | **Source** |
| --- | --- | --- | --- | --- | --- |
| CD16 | eBioCD16 | 1-9161-71 | 4304474 | FITC | Ebioscience |
| CD3 | OKT3 | 317308 | B256076 | PE | Biolegend |
| CD56 | HCD56 | 318306 | B252053 | PE | Biolegend |
| CD19 | HIB19 | 302254 | B227178 | PE | Biolegend |
| CCR2 | K036C2 | 357212 | B260108 | PE/Cy7 | Biolegend |
| ICAM | HCD54 | 322718 | B193832 | AF 647 | Biolegend |
| CD45 | 2D1 | 368514 | B248834 | AF 700 | Biolegend |
| CD14 | M5E2 | 301820 | B274258 | APC/Cy7 | Biolegend |
| HLA-DR | L243 | 307624 | B278326 | Pacific Blue | Biolegend |
| CD66b | G10F5 | 305106 | B278603 | PE | Biolegend |
| SiglecF | E50-2440 | 552126 | 7058859 | PE | BD<br>Biosciences |
| CD11b | M1/70 | 101256 | B238075 | PE Dazzle | BD<br>Biosciences |
| CD11b | M1/70 | 101243 | B253527 | BV785 | Biolegend |
| MHCII | M5.114.15.2 | 107624 | B267551 | PerCP | Ebioscience |
| Epcam | G8.8 | 118230 | B251914 | APCCy7 Fire | Biolegend |
| CD3 | 17A2 | 100244 | B198733 | BIOTIN | Biolegend |
| CD3 | 17A2 | 100213 | B261416 | Pacific Blue | Biolegend |
| CD3 | 17A2 | 100229 | B282101 | BV 650 | Biolegend |
| CD3 | 17A2 | 100206 | B210714 | PE | Biolegend |
| CD19 | 6D5 | 115541 | B242632 | BV650 | Biolegend |
| CD19 | 6D5 | 115504 | B244881 | Biotin | Biolegend |
| CD19 | 6D5 | 115526 | B265435 | Pacific Blue | Biolegend |
| CD19 | 6D5 | 115508 | B223615 | PE | Biolegend |
| CD103 | 2E7 | 121433 |  | BV605 | Biolegend |
| Ly6G | 1A8 | 127604 | B218526 | BIOTIN | Biolegend |
| Ly6G | 1A8 | 127608 | B221647 | PE | Biolegend |
| Ly6G | 1A8 | 127628 | B280589 | BV 421 | Biolegend |
| Ly6G | 1A8 | 135512 | B213676 | AF 488 | Biolegend |
| Lyve-1 | ALY7 | 50-0443-82 |  | eFluor 660 | Ebioscience |
| CD115 | AFS98 | 135510 | B211309 | APC | Biolegend |
| CD115 | AFS98 | 128006 | B217035 | FITC | Biolegend |
| Ly6C | HK1.4 | 128032 | B232012 | BV 421 | Biolegend |
| Pan-CD45 | 30-F11 | 103128 | B274307 | AF 700 | Biolegend |
| CD11c | N418 | 117318 | B222652 | PE/Cy7 | Biolegend |
| CD11c | N418 | 117352 | B218048 | APC/Fire750 | Biolegend |
| CD64 | X54-5/7.1 | 139304 | B191540 | PE | Biolegend |
| CD64 | X54-5/7.1 | 139306 | B207411 | APC | Biolegend |
| CD4 | H129.19 | 553649 |  | Biotin | BD<br>Biosciences |
| CD5 | 53-7.3 | 553019 |  | Biotin | BD<br>Biosciences |
| CD8a | 53 -6.7 | 553029 |  | Biotin | BD<br>Biosciences |
| CD11b | M1/70 | 101256 | B238075 | Biotin | BD<br>Biosciences |
| CD45R/B220 | RA3-6B2 | 553086 |  | Biotin | BD<br>Biosciences |
| Ter119 | TER-119 | 553672 |  | Biotin | BD<br>Biosciences |
| Gr-1/Ly-6G/C | RB6-8C5 | 553125 |  | Biotin | BD<br>Biosciences |
| CD117/cKit | 2B8 | 105811 | B249345 | APC | Biolegend |
| Sca-1/Ly- | E13-161.7 | 122506 |  | FITC | Biolegend |
| CD48 | HM48-1 | 103406 |  | PE | Biolegend |
| CD150 | 12F12.2 | 115914 |  | PECy7 | Biolegend |
| CD71 | RI7217 | 113807 |  | PE | Biolegend |
| Streptavidin | - | 405232 | B251688 | BV 650 | Biolegend |
| Streptavidin | - |  |  | Pacific Blue | BD<br>biosciences |
| LIVE/DEAD®<br>Fixable Aqua | - | L34957 | 2068285 | UV650 | Life<br>Technologies<br>or Biolegend |
| CD45.2 | 104 | 109822 | B252126 | AF700 | Biolegend |
| CD45.1 | A20 | 110741 | B253101 | BV510 | Biolegend |
| Sca.1 | D7 | 108129 | B262926 | BV510 | Biolegend |
| CD150 | <u>TC15-</u><br><u>12F12.2</u> | 115903 |  | PE | Biolegend |
| CD105 | MJ7/18 | 120412 | B245562 | PacBlue | Biolegend |
| CD41 | MWReg30 | 133927 | B268849 | APCCy7 | Biolegend |
| IFNAR | <u>MAR1-5A3</u> | 127325 | B286788 | PECy7 | Biolegend |
| CD5 | 53-7.3 | 100603 | B254317 | Biotin | Biolegend |
| Ly6G | 1A8 | 127604 | B218529 | Biotin | Biolegend |
| B220 | RA3-6B2 | 103204 | B288658 | Biotin | Biolegend |
| CD11b | M1/70 | 562287 |  | CF594 | BD<br>Biosciences |

#### Gating strategies

Human monocytes: Singles Dapi^-^CD45^+^non-granulocyte Lin(CD3/CD56/CD19/CD66b)^-^ HLADR^+^ CD14^+^ and/or CD16^+^ cells

Mouse blood monocytes: Singles Dapi^-^CD45^+^ Lin(CD3/CD19/ Ly6G)^-^ CD115^+^CD11b^+^Ly6C^hi^, Ly6C^int^ or Ly6C^-^

Mouse blood neutrophils: Singles, Dapi^-^CD45^+^Ly6G^+^CD11b^+^Ly6C ^int^

Mouse lung/ BAL alveolar macrophages: Singles, Zombie Aqua^-^CD45^+^Lin (CD3/CD19/Ly6G)^-^CD64^+^SiglecF^+^CD11c^+^

Mouse lung interstitial / BAL inflammatory macrophages: Singles, Zombie Aqua^-^CD45^+^Lin (CD3/CD19/Ly6G)^-^CD64^+^SiglecF^-^CD11c^+/-^ then Ly6C^+/-^ MHCII^+/-^

Lung monocytes: Singles, Zombie Aqua^-^SinglesCD45^+^Lin (CD3/CD19/Ly6G)^-^CD64^int^ CD11b^+^ Ly6C^+^

Lung/ BAL neutrophils: Singles Aqua or Dapi^-^, CD45^+^CD11b^+^Ly6G^+^ Lung cDC1 Zombie Aqua-Singles CD45^+^CD11c^hi^,CD103^+^, CD64^-^MHCII^+^

BM HSPC SLAM analysis Alive, Singles LK (Lin-cKit^+^) and LSK (Lin-cKit^+^Sca-1^+^) cells. LSK cells were further sub-gated on hematopoietic stem cells (HSCs: LSK CD48- CD150^+^), multipotent progenitors (MPPs: LSK CD48^-^CD150^-^), hematopoietic progenitor cells-1 (HPC-1: LSK CD48^+^CD150^-^) and hematopoietic progenitor cells-2 (HPC-2: LSK CD48^+^CD150^+^)

BM erythroid progenitors based on Pronk analysis ^26^ Singles, Dapi or Aqua^-^, Lin^-^, CD11b^-^ , cKit^+^, Sca1^-^, CD32/16^-^, CD41^-^, CD105^+^ or CD150^+^ (Pre-MegE CD150^+^CD105^-^, Pre- CFUE CD150^+^CD105^+^, CFUE CD150^-^CD105^+^)

Further gating strategy information can be made available upon request.

#### BAL/ serum Cytokine/ Chemokine quantification

BAL and serum supernatants were collected and stored at -80°C until use. Cytokine and chemokine levels were measured using an MSD V-plex plate as per manufacturer’s instructions.

#### Lung injury measurements

IgM BAL levels were measured using Ab133047 Abcam kit as per manufacturer’s instructions.

BAL LDH activity (measured as colorimetric reduction of NAD to NADH) was performed using Ab102526 (Abcam) as per manufacturer’s instructions.

BAL total protein was measured using Pierce BCA Assay (Thermofisher) as per manufacturer’s instructions

#### In vitro bone marrow culture

Naïve wild-type C57BL/6 bone marrow was obtained flushing the femoral and tibial bones and RBC were lysed. Cell were cultured in hypoxia (FiO2 1%) or normoxia (FiO2 21%) with conditioned DMEM for 1 hour prior to addition of IFNb 10ng/ml (RnD 8234-MB-010) for a further 3 hours. Cell pellets were collected and Qiagen RLT buffer added (containing 10uL/ml ß-mercaptoethanol). Pellets were snap frozen and stored at -80 for RNA extraction.

#### RNA isolation and relative quantification

RNA was isolated from bone marrow cells using the gDNA eliminator solution for purification of total RNA (RNeasy Plus Mini Kit, Qiagen). cDNA was synthesized by using AMV reverse transcriptase with random primers (Promega). TaqMan gene expression assays (Applied Biosystems, Thermo Fisher) and PrimeTime qPCR Probe Assays (IDT) were used for relative quantification of cDNA using SDS 2.4 (Thermo Fisher) and normalized to ACTB expression.

#### Immunohistochemistry

Murine paraffin-embedded blocks were prepared from lungs fixed via the trachea with 10% buffered formalin. The lung sections were stained with anti-IL10 (ab189392, Abcam), anti-F4/80 (ab6640, Abcam) or isotype control after deparaffinization and antigen retrieval. The following were used TSA plus system amplification (NEL744B001KT, Perkin Elmer) and autofluorescence quenching with TrueView (Vector, SP-8400). The nuclei were stained with DAPI (422801, Sigma-Aldrich). Images were obtained using inverted Di8 microscope (Carl Zeiss).

#### nCounter NanoString platform analysis

For human monocytes 5000 cells were sorted using the aforementioned human monocyte gating strategy directly into 2ul RLT buffer using a BD Fusion Sorter. 5000 mouse classical monocytes were sorted from mice treated with LPS and housed in normoxia, hypoxia and hypoxia+CSF1 gating on single DAPI^-^CD45^+^Lin^-^CD115^+^Ly6C^hi^ cells into 2ul RLT. Cell pellets were vortexed and centrifuged prior to immediate freezing until ready for processing. Human and mouse myeloid inflammation NanoString gene expression plates were run as per manufacturer’s instructions at the University of Edinburgh HTPU Centre within the MRC Institute of Genetics and Molecular Medicine/Cancer Research UK Edinburgh Centre.

#### Proteomic analysis

Sorted monocytes were processed for proteomics using the ‘in cell digest”, as described by Kelly et al. bioRxiv 2020^52^, resuspended in digestion buffer (0.1 M triethylammonium bicarbonate + 1 mM MgCl2) and digested with benzonase (>99%, Millipore) for 30 min at 37 °C, followed by trypsin (Thermo Fisher Scientific, 1:50 w/w protein) overnight at 37 °C. A second aliquot of trypsin (1:50) was subsequently added and incubated at 37 °C for 4 hours. A minimum of 25 ng of trypsin was added. Digests were acidified and desalted using StageTips^53^ and either subjected to tip-based fractionation or direct analysis by LC- MS/MS.

Following digestion, and in order to generate the reference spectral library, peptides were subjected to reverse phase high pH tip fractionation following the general guidelines as described by Rappsilber et al^53^. In brief, tips for fractionation were made using three SDB- XC disks (Merck, UK) per tip. The tip was cleaned and conditioned using sequentially methanol, 80% acetonitrile (MeCN) (Thermo Fisher Scientific, UK) in 0.1% NH4OH (v/v) and 0.1% NH4OH (52mM) (v/v). Peptides, resuspended also 0.1% NH4OH (pH=10), were spun through the SDB-XC disks and the flow-through was collected, acidified and concentrated on C-18 stage-Tips before subjected to MS analysis. Fractionation was then achieved by sequential elution with 7%, 14%, 21%, 28%, 35%, 55%, and 80% MeCN in 0.1% NH4OH. Fractions were then dried at ambient temperature (Concentrator 5301, Eppendorf, UK) and prepared for MS analysis by resuspension in 6uL of 0.1% TFA.

Data-dependent acquisition (DDA) LC-MS analyses were performed on an Orbitrap Fusion™ Lumos™ Tribrid™ Mass Spectrometer (Thermo Fisher Scientific, UK) coupled on-line, to an Ultimate 3000 HPLC (Dionex, Thermo Fisher Scientific, UK). Peptides were separated on a 50 cm (2 µm particle size) EASY-Spray column (Thermo Scientific, UK), which was assembled on an EASY-Spray source (Thermo Scientific, UK) and operated constantly at 50°C. Mobile phase A consisted of 0.1% formic acid in LC-MS grade water and mobile phase B consisted of 80% acetonitrile and 0.1% formic acid. Peptides were loaded onto the column at a flow rate of 0.3 μL min^-1^ and eluted at a flow rate of 0.25 μL min^-1^ according to the following gradient: 2 to 40% mobile phase B in 120 min and then to 95% in 11 min. Mobile phase B was retained at 95% for 5 min and returned back to 2% a minute after until the end of the run (160 min in total).

The spray voltage was set at 2.2kV and the ion capillary temperature at 280°C. Survey scans were recorded at 60,000 resolution (scan range 400-1600 m/z) with an ion target of 1.0E6, and injection time of 50ms. MS2 was performed in the orbitrap (resolution at 15,000), with ion target of 5.0E4 and HCD fragmentation^54^ with normalized collision energy of 27. The isolation window in the quadrupole was 1.4 Thomson. Only ions with charge between 2 and 6 were selected for MS2. Dynamic exclusion was set at 60 s. The cycle time was set at 3 seconds.

Samples subjected to Data Independent Acquisition (DIA), were prepared for MS analysis by resuspension in 0.1% TFA. MS Analyses were performed on an Orbitrap Fusion™ Lumos™ Tribrid™ Mass Spectrometer (Thermo Fisher Scientific, UK). LC conditions (instrumentation, column, and gradient) were the same as described above.

Survey scans were performed at 15,000 resolution, with scan range at 350-1500 m/z, maximum injection time at 50ms and AGC target at 4.5E5. MS/MS DIA was performed in the orbitrap at 30,000 resolution with a scan range of 200-2000 m/z. The mass range was set to “normal” the maximum injection time to 54ms and the AGC target to 2.0E5. The inclusion mass list with the correspondent isolation windows are shown in the table below. Data for both survey and MS/MS scans were acquired in profile mode. A blank sample (0.1% TFA, 80% MeCN 1:1 v/v) was run in between of each sample to avoid carryover.

| m/z | z | t start<br>(min) | t stop<br>(min) | Isolation<br>Window<br>(m/z) |
| --- | --- | --- | --- | --- |
| 410 | 3 | 0 | 155 | 20 |
| 430 | 3 | 0 | 155 | 20 |
| 450 | 3 | 0 | 155 | 20 |
| 470 | 3 | 0 | 155 | 20 |
| 490 | 3 | 0 | 155 | 20 |
| 510 | 3 | 0 | 155 | 20 |
| 530 | 3 | 0 | 155 | 20 |
| 550 | 3 | 0 | 155 | 20 |
| 570 | 3 | 0 | 155 | 20 |
| 590 | 3 | 0 | 155 | 20 |
| 610 | 3 | 0 | 155 | 20 |
| 630 | 3 | 0 | 155 | 20 |
| 650 | 3 | 0 | 155 | 20 |
| 670 | 3 | 0 | 155 | 20 |
| 690 | 3 | 0 | 155 | 20 |
| 710 | 3 | 0 | 155 | 20 |
| 730 | 3 | 0 | 155 | 20 |
| 750 | 3 | 0 | 155 | 20 |
| 770 | 3 | 0 | 155 | 20 |
| 790 | 3 | 0 | 155 | 20 |
| 820 | 3 | 0 | 155 | 40 |
| 860 | 3 | 0 | 155 | 40 |
| 910 | 3 | 0 | 155 | 60 |
| 970 | 3 | 0 | 155 | 60 |

### QUANTIFICATION AND STATISTICAL ANALYSIS

#### Gene expression analysis

Normalization of data was carried out using the geNorm selection of housekeeping genes function on NanoString nCounter analysis software. Resulting Log2 normalized values were used in subsequent analyses. Differential genes (“DE genes) were defined as genes with log2 FC>1, P-value <0.05 across sample groups. Hierarchical clustering of sets of DE genes was carried out using Euclidian and Ward methods based on Pearson correlation values across transcriptional scores. Z-score scalar normalization of data was applied to the data prior to plotting as heatmaps. Analyses, including the drawing of heatmaps and volcano plots were carried out in R using the package ggplot2 (https://cran.r-project.org/web/packages/ggplot2/index.html). Analysis of datasets was carried out by Thomson Bioinformatics, Edinburgh, UK.

#### Proteomic analysis

MS raw data files were processed using Spectronaut v14.7.201007.47784 using a human reference FASTA sequence from UniProt using default search parameters. The resulting protein-level data were analysed using R v3.5.0. Protein parts-per-million (ppm) intensities were calculated by dividing the abundance of a protein by the total abundance measured in the sample and multiplied by a million. Fold-changes were calculated by dividing the mean ppm intensities between ARDS and healthy control samples. P-values were calculated using a t-test on log-transformed ppm intensities. Proteins were designated as significantly changing if they showed p-values < 0.05 and fold changes exceeding 1.96 standard deviations away from the mean (i.e., z-score > 1.96). Only proteins that were quantitated in all samples are shown in the volcano plot.

#### Quantification and statistical analysis

Statistical tests were performed using Prism 8.00 and 9.0.2 software (GraphPad Software Inc) (specific tests detailed in figure legends). Significance was defined as a p value of <0.05 (after correction for multiple comparisons where applicable). Sample sizes (with each n number representing a different blood donor for human cells or an individual mouse for animal experiments) are shown in figure legends.

## References

1. Bellani, G., et al. Epidemiology, Patterns of Care, and Mortality for Patients With Acute Respiratory Distress Syndrome in Intensive Care Units in 50 Countries. JAMA 315, 788–800 (2016).

2. Fan, E. et al. An Official American Thoracic Society/European Society of Intensive Care Medicine/Society of Critical Care Medicine Clinical Practice Guideline: Mechanical Ventilation in Adult Patients with Acute Respiratory Distress Syndrome. Am J Respir Crit Care Med 195, 1253–1263 (2017).

3. Steinberg, K.P. et al. Evolution of bronchoalveolar cell populations in the adult respiratory distress syndrome. Am J Respir Crit Care Med 150, 113–122 (1994).

4. Park, W.Y. et al. Cytokine balance in the lungs of patients with acute respiratory distress syndrome. Am J Respir Crit Care Med 164, 1896–1903 (2001).

5. Tarling, J.D., Lin, H.S. & Hsu, S. Self-renewal of pulmonary alveolar macrophages: evidence from radiation chimera studies. J Leukoc Biol 42, 443–446 (1987).

6. Murphy, J., Summer, R., Wilson, A.A., Kotton, D.N. & Fine, A. The prolonged life-span of alveolar macrophages. Am J Respir Cell Mol Biol 38, 380–385 (2008).

7. Guilliams, M. et al. Alveolar macrophages develop from fetal monocytes that differentiate into long-lived cells in the first week of life via GM-CSF. J Exp Med 210, 1977–1992 (2013).

8. Hashimoto, D. et al. Tissue-resident macrophages self-maintain locally throughout adult life with minimal contribution from circulating monocytes. Immunity 38, 792–804 (2013).

9. Bedoret, D. et al. Lung interstitial macrophages alter dendritic cell functions to prevent airway allergy in mice. J Clin Invest 119, 3723–3738 (2009).

10. Sabatel, C. et al. Exposure to Bacterial CpG DNA Protects from Airway Allergic Inflammation by Expanding Regulatory Lung Interstitial Macrophages. Immunity 46, 457–473 (2017).

11. Zhou, B. et al. The angiocrine Rspondin3 instructs interstitial macrophage transition via metabolic-epigenetic reprogramming and resolves inflammatory injury. Nat Immunol (2020).

12. Chakarov, S. et al. Two distinct interstitial macrophage populations coexist across tissues in specific subtissular niches. Science 363 (2019).

13. Patel, A.A. et al. The fate and lifespan of human monocyte subsets in steady state and systemic inflammation. J Exp Med 214, 1913–1923 (2017).

14. Xu, H., Manivannan, A., Crane, I., Dawson, R. & Liversidge, J. Critical but divergent roles for CD62L and CD44 in directing blood monocyte trafficking in vivo during inflammation. Blood 112, 1166–1174 (2008).

15. Schenkel, A.R., Mamdouh, Z., Chen, X., Liebman, R.M. & Muller, W.A. CD99 plays a major role in the migration of monocytes through endothelial junctions. Nat Immunol 3, 143–150 (2002).

16. Zhao, Q. et al. The role of mitogen-activated protein kinase phosphatase-1 in the response of alveolar macrophages to lipopolysaccharide: attenuation of proinflammatory cytokine biosynthesis via feedback control of p38. J Biol Chem 280, 8101–8108 (2005).

17. Bhattacharyya, S., Borthakur, A., Dudeja, P.K. & Tobacman, J.K. Lipopolysaccharide-induced activation of NF-kappaB non-canonical pathway requires BCL10 serine 138 and NIK phosphorylations. Exp Cell Res 316, 3317–3327 (2010).

18. Mimura, I. et al. Dynamic change of chromatin conformation in response to hypoxia enhances the expression of GLUT3 (SLC2A3) by cooperative interaction of hypoxia- inducible factor 1 and KDM3A. Mol Cell Biol 32, 3018–3032 (2012).

19. Chen, W.K. et al. CREB Negatively Regulates IGF2R Gene Expression and Downstream Pathways to Inhibit Hypoxia-Induced H9c2 Cardiomyoblast Cell Death. Int J Mol Sci 16, 27921–27930 (2015).

20. Abu-El-Rub, E. et al. Hypoxia-induced shift in the phenotype of proteasome from 26S toward immunoproteasome triggers loss of immunoprivilege of mesenchymal stem cells. Cell Death Dis 11, 419 (2020).

21. Sammarco, M.C., Ditch, S., Banerjee, A. & Grabczyk, E. Ferritin L and H subunits are differentially regulated on a post-transcriptional level. J Biol Chem 283, 4578–4587 (2008).

22. Gray, L.H. & Steadman, J.M. Determination of the Oxyhaemoglobin Dissociation Curves for Mouse and Rat Blood. J Physiol 175, 161–171 (1964).

23. Aggarwal, N.R., King, L.S. & D’Alessio, F.R. Diverse macrophage populations mediate acute lung inflammation and resolution. Am J Physiol Lung Cell Mol Physiol 306, L709–725 (2014).

24. Meduri, G.U. et al. Persistent elevation of inflammatory cytokines predicts a poor outcome in ARDS. Plasma IL-1 beta and IL-6 levels are consistent and efficient predictors of outcome over time. Chest 107, 1062–1073 (1995).

25. Bain, C.C. et al. Resident and pro-inflammatory macrophages in the colon represent alternative context-dependent fates of the same Ly6Chi monocyte precursors. Mucosal Immunol 6, 498–510 (2013).

26. Pronk, C.J. et al. Elucidation of the phenotypic, functional, and molecular topography of a myeloerythroid progenitor cell hierarchy. Cell Stem Cell 1, 428–442 (2007).

27. Lucas, C. et al. Longitudinal analyses reveal immunological misfiring in severe COVID-19. Nature 584, 463–469 (2020).

28. Lasseaux, C., Fourmaux, M.P., Chamaillard, M. & Poulin, L.F. Type I interferons drive inflammasome-independent emergency monocytopoiesis during endotoxemia. Sci Rep 7, 16935 (2017).

29. Silver, R.T. Recombinant interferon-alpha for treatment of polycythaemia vera. Lancet 2, 403 (1988).

30. Yoshida, H., Okabe, Y., Kawane, K., Fukuyama, H. & Nagata, S. Lethal anemia caused by interferon-beta produced in mouse embryos carrying undigested DNA. Nat Immunol 6, 49–56 (2005).

31. Gow, D.J. et al. Characterisation of a novel Fc conjugate of macrophage colony-stimulating factor. Mol Ther 22, 1580–1592 (2014).

32. Bain, C.C. et al. Long-lived self-renewing bone marrow-derived macrophages displace embryo-derived cells to inhabit adult serous cavities. Nat Commun 7, ncomms11852 (2016).

33. Sheikh, M.H. & Solito, E. Annexin A1: Uncovering the Many Talents of an Old Protein. Int J Mol Sci 19 (2018).

34. Proto, J.D. et al. Regulatory T Cells Promote Macrophage Efferocytosis during Inflammation Resolution. Immunity 49, 666–677 e666 (2018).

35. Rosseau, S. et al. Phenotypic characterization of alveolar monocyte recruitment in acute respiratory distress syndrome. Am J Physiol Lung Cell Mol Physiol 279, L25–35 (2000).

36. Abraham, E. Neutrophils and acute lung injury. Crit Care Med 31, S195–199 (2003).

37. Thompson, A.A. et al. Hypoxia determines survival outcomes of bacterial infection through HIF-1alpha dependent re-programming of leukocyte metabolism. Sci Immunol 2 (2017).

38. Larson, S.R. et al. Ly6C(+) monocyte efferocytosis and cross-presentation of cell-associated antigens. Cell Death Differ 23, 997–1003 (2016).

39. Soehnlein, O. & Lindbom, L. Phagocyte partnership during the onset and resolution of inflammation. Nat Rev Immunol 10, 427–439 (2010).

40. Erslev, A.J. Erythroid adaptation to altitude. Blood Cells 7, 495–508 (1981).

41. Hannon, J.P., Shields, J.L. & Harris, C.W. Effects of altitude acclimatization on blood composition of women. J Appl Physiol 26, 540–547 (1969).

42. Xie, J. et al. Association Between Hypoxemia and Mortality in Patients With COVID-19. Mayo Clin Proc 95, 1138–1147 (2020).

43. Makarova, K. et al. Comparative genomics of the lactic acid bacteria. Proc Natl Acad Sci U S A 103, 15611–15616 (2006).

44. Yanez, A. et al. Granulocyte-Monocyte Progenitors and Monocyte-Dendritic Cell Progenitors Independently Produce Functionally Distinct Monocytes. Immunity 47, 890–902 e894 (2017).

45. Dunning, J. et al. Progression of whole-blood transcriptional signatures from interferon-induced to neutrophil-associated patterns in severe influenza. Nat Immunol 19, 625–635 (2018).

46. Seo, S.U. et al. Type I interferon signaling regulates Ly6C(hi) monocytes and neutrophils during acute viral pneumonia in mice. PLoS Pathog 7, e1001304 (2011).

47. Kashani, K.B. Hypoxia in COVID-19: Sign of Severity or Cause for Poor Outcomes. Mayo Clin Proc 95, 1094–1096 (2020).

48. Liao, M. et al. Single-cell landscape of bronchoalveolar immune cells in patients with COVID-19. Nat Med 26, 842–844 (2020).

49. Guilliams, M., Thierry, G.R., Bonnardel, J. & Bajenoff, M. Establishment and Maintenance of the Macrophage Niche. Immunity 52, 434–451 (2020).

50. Lim, H.Y. et al. Hyaluronan Receptor LYVE-1-Expressing Macrophages Maintain Arterial Tone through Hyaluronan-Mediated Regulation of Smooth Muscle Cell Collagen. Immunity 49, 1191 (2018).

51. Alivernini, S. et al. Distinct synovial tissue macrophage subsets regulate inflammation and remission in rheumatoid arthritis. Nat Med 26, 1295–1306 (2020).

52. Kelly, V., al-Rawi, A., Lewis, D. & Ly, T. Cell cycle state proteomics and classification using in-cell protease digests and mass spectrometry. bioRxiv, 2020.2007.2003.186023 (2020).

53. Rappsilber, J., Mann, M. & Ishihama, Y. Protocol for micro-purification, enrichment, pre-fractionation and storage of peptides for proteomics using StageTips. Nat Protoc 2, 1896–1906 (2007).

54. Olsen, J.V. et al. Higher-energy C-trap dissociation for peptide modification analysis. Nat Methods 4, 709–712 (2007).

