## Supplementary data for "Hypoxia shapes the immune landscape in lung injury promoting inflammation persistence"

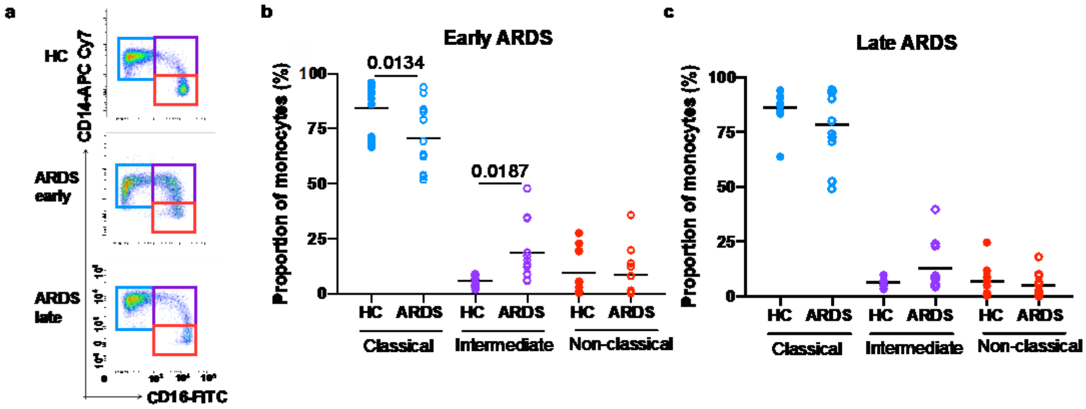


**Figure S1. Monocyte sub-populations are altered early in ARDS**

(**a**) Representative plots and proportions of monocyte sub-populations based on CD14 and CD16 expression early and (**b**) late (**c**). **b**, **c** one-way ANOVA with Holm-Sidak post-test.

a

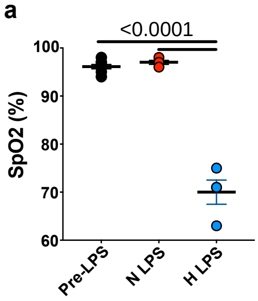


**Figure S2. Oxygen saturations in mice post LPS injury** (**a**) Oxygen saturations in mice were measured in mice at baseline pre-LPS nebulisation (pre-LPS) and 6 hours post-LPS (N LPS- mice housed in normoxia post-LPS, H LPS- mice housed in hypoxia post-LPS). (Pre-LPS n=6, N LPS n=3, H LPS n=3). Statistical testing performed using one-way ANOVA with Tukey's multiple comparisons test.

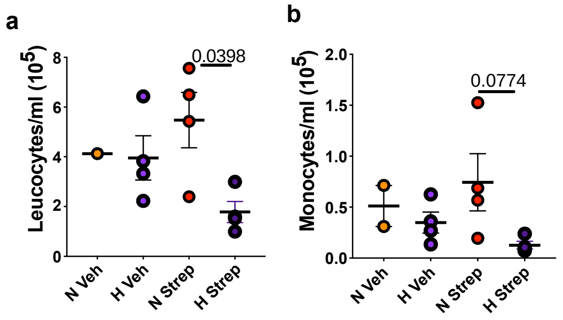


**Figure S3. *Streptococcus pneumoniae* infection in hypoxia leads to leukopenia and monocytopenia.** Mice were inoculated with *Streptococcus pneumoniae* (Strep) or vehicle (Veh) intratracheally (i.t.) and housed in normoxia (N) or hypoxia (H) until 24 hours post-i.t. (**a**) Blood cell counts and (**b**) monocyte counts mice housed in normoxia (N) or hypoxia (H) for 24 hours (Veh N and H n=4, Strep N and H n=6). Statistical testing performed using one-way ANOVA with Tukey's multiple comparisons test.

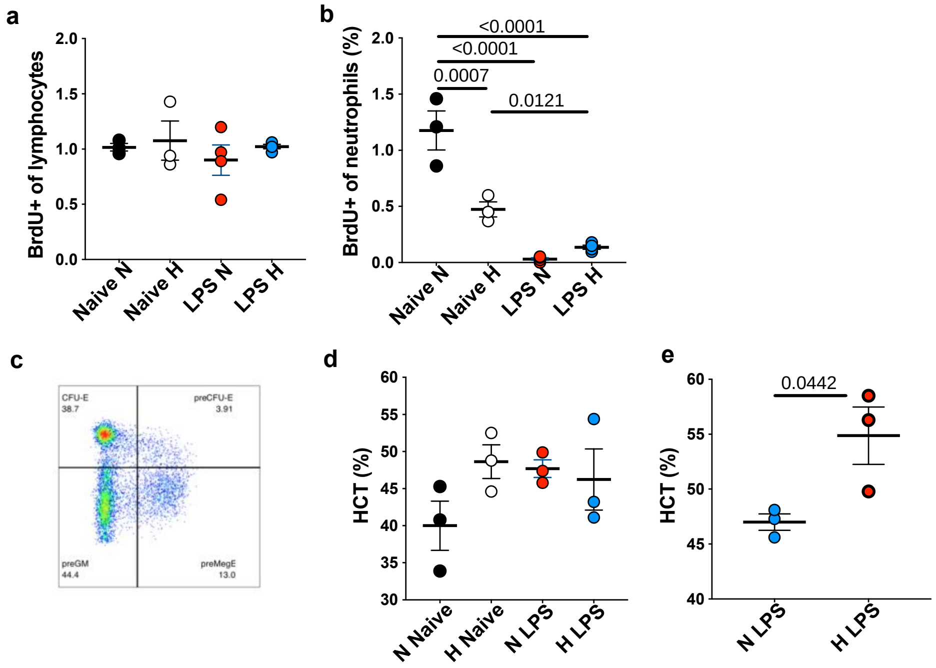


**Figure S4. Impact of hypoxia on bone marrow cell egress and composition** (**a**) BrdU+ blood lymphocytes (CD3 and CD19$^+$) and (**b**) BrdU+ blood neutrophil proportion in mice treated with LPS and housed in normoxia (N) and hypoxia (H) for 24 hours (Naïve N and H n=3, LPS N and H n=4). Data representative of 2 experiments (**c**) gating strategy for bone marrow common myeloid progenitor progeny on CD41$^-$ CD16/32$^-$ cells. (**d**) blood hematocrit at 24 hours (n=3/ group) data representative of 3 experiments or (**e**) 5 days in mice treated with LPS and housed in normoxia (N) and hypoxia (H) (n=3/group). Data representative of 2 experiments (**a**, **b**) Statistical testing performed using one-way ANOVA with Tukey's multiple comparisons test, (**e**) statistical testing performed using two-tailed unpaired t-test.

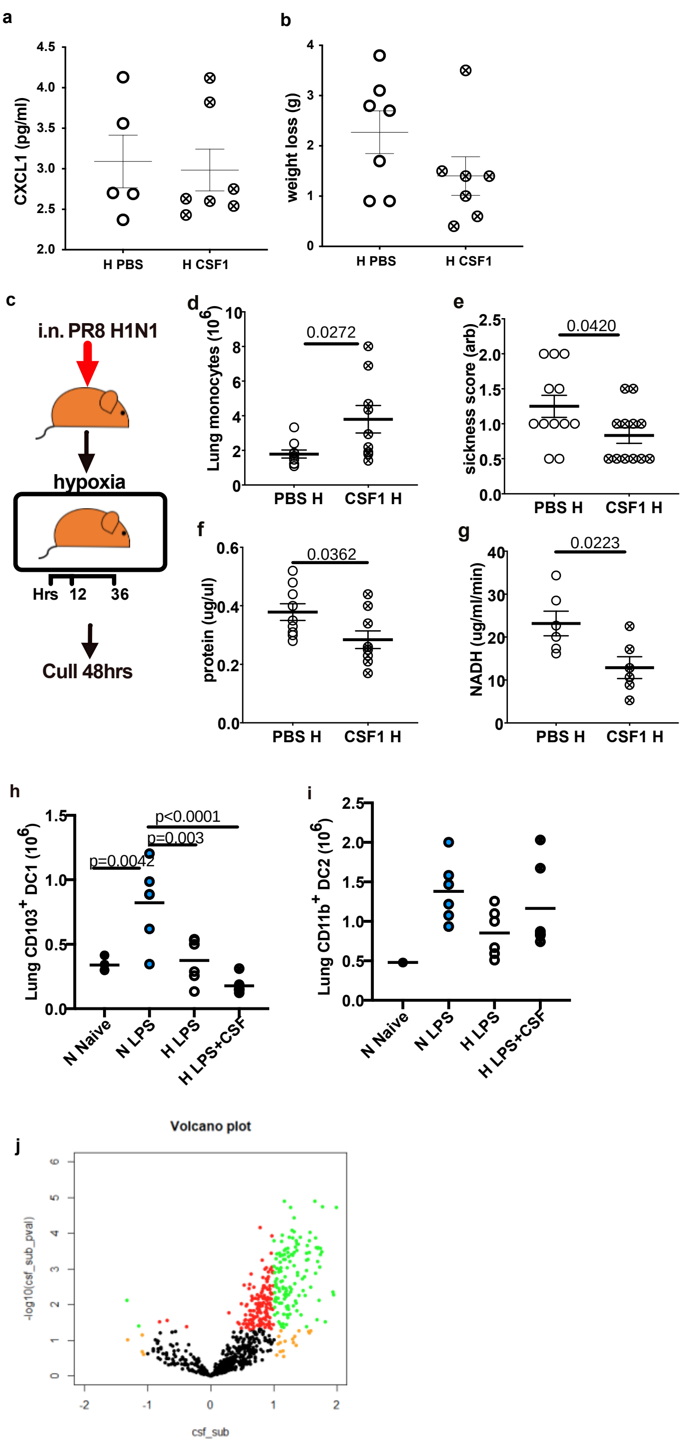


**Figure S5. CSF1-Fc alters monocyte phenotype and improves injury outcomes**

(**a**) BAL CXCL1 (**b**) weight loss in hypoxic LPS-induced ALI treated with PBS (H PBS) or CSF1 (H CSF1)(**c**) Schematic of virally-induced ALI plaque-forming units (p.f.u.).

(**d**) Lung monocyte numbers, (**e**) arbitrary sickness scores, (**f**) BAL protein and (**g**) LDH activity (as measured by NADH) were measured at 48 hours in mice with virally-induced ALI housed in hypoxia and receiving either PBS or CSF1. **(h**) Lung cDC1 (gated on Alive $CD45^{+}$Lin-$CD64^{-}CD11c^{+}Cd103^{+}$) and (**i**) cDC2 (gated on Alive $CD45^{+}$Lin-$CD64^{-}CD11c^{+}Cd103^{-}$ $CD11b^{+}$) chimerism relative to blood monocyte chimerism in non-LPS-treated (naïve) or LPS-treated mice housed in normoxia (N) or hypoxia (H) and receiving PBS or CSF1. (**j**) Volcano plot of measured genes in classical monocytes from mice treated with LPS and housed in hypoxia for 5 days and treated with PBS or CSF1-Fc for 4 days. Green dots correspond to genes with log2 change > 1 or < -1 and with p values <0.05. Orange dots denote genes changing > 1 or < -1 with p values >0.05. Red dots denote genes with log2 changes out-with the differential expression range but with p values <0.05. Data shown as mean ±SEM with each data point representing an individual mouse. Statistics **d** two-tailed Mann-Whitney following D'Agostino & Pearson normality test, **f**, **g** One way ANOVA with Tukeys' multiple comparisons test, **h, I**iOne way ANOVA with Kruskal-Wallis multiple comparisons test.

### Table S1. Patient demographics, clinical severity and oxygenation

| Group | Gender | Age | APACHE II score | Lowest recorded PaO2 prior to sampling | FiO2(%) at time of pre-sample oxygenation recording | Cause of ARDS: Pulmonary or Extrapulmonary |
|---|---|---|---|---|---|---|
| Early | F | 58 | 19 | 9.8 | 70 | Extrapulmonary |
| Early | M | 78 | 31 | 8.1 | 100 | Extrapulmonary |
| Early | M | 73 | 19 | 9.1 | 21 | Pulmonary |
| Early | F | 46 | 15 | 10 | 100 | Pulmonary |
| Late | M | 72 | 21 | 7.9 | 40 | Extrapulmonary |
| Late | F | 72 | 17 | 9.1 | 35 | Extrapulmonary |
| Late | M | 74 | 9 | 8.9 | 40 | Extrapulmonary |
| Late | F | 53 | 30 | 6.9 | 45 | Extrapulmonary |
| Early | M | 63 | 19 | 9 | 25 | Pulmonary |
| Late | M | 49 | 16 | 7 | 45 | Pulmonary |
| Early | F | 72 | 29 | 7.3 | 80 | Pulmonary |
| Late | M | 57 | 38 | 8.4 | 60 | Pulmonary |
| Early | F | 51 | 24 | 7.8 | 30 | Pulmonary |
| Early | M | 67 | 21 | 9.2 | 50 | Pulmonary |
| Early | F | 42 | 21 | 12 | 70 | Extrapulmonary |
| Early | M | 54 | 7 | 6.1 | 70 | COVID+ ARDS |
| Late | F | 41 | 13 | 9.7 | 60 | COVID+ ARDS |
| Late | M | 32 | 21 | 14.1 | 55 | Extrapulmonary |
| Early | M | 54 | 27 | 8.4 | 30 | Pulmonary |
| Early | F | 56 | 13 | 9.5 | 45 | COVID+ ARDS |
| Late | F | 60 | 13 | 7.7 | 35 | Extrapulmonary |
| Late | M | 53 | 18 | 15 | 75 | Pulmonary |
| Late | M | 53 | 11 | 8.2 | 60 | Pulmonary |
